# Bacterial Aminopeptidase–Activated Peptide Prodrug Enables Species-Selective Targeting of *Pseudomonas aeruginosa*

**DOI:** 10.64898/2026.03.29.715093

**Authors:** Qingtian Gong, Silvia Synowsky, Aisling Lynch, John R. F. B. Connolly, Neeladri Sekhar Roy, Sally L. Shirran, Megan Bergkessel, Marc Devocelle, Clarissa M. Czekster

**Affiliations:** University of St Andrews, School of Biology, Biomolecular Sciences building, North Haugh, KY16 9ST, UK; University of St Andrews, BSRC Mass Spectrometry and Proteomics Facility, Schools of Biology and Chemistry, North Haugh, St Andrews, KY16 9ST, UK; RCSI University of Medicine and Health Sciences, Department of Chemistry, 123, St. Stephen’s Green, D02 YN77, Dublin 2, Ireland; The Group of Applied Physics, School of Physics, Clinical and Optometric Sciences, Technological University Dublin, D07 ADY7 Dublin, Ireland; School of Life Sciences, University of Dundee, Dundee, UK

## Abstract

*Pseudomonas aeruginosa* is an adaptable organism, frequently found in chronic infections, and for which antimicrobial resistance is a growing concern. Therefore, there is an urgent need for alternative therapeutic strategies. Cationic antimicrobial peptides (AMPs) offer potent bactericidal activity but suffer from limited selectivity and potential host toxicity. To enhance species-specific targeting, we designed two prodrug variants of the AMP D-Bac8C^Leu2,5^ – EEEE□-Bac8C^Leu2,5^ and ELE□ D-Bac8C^Leu2,5^ —engineered for activation by the *P. aeruginosa* extracellular aminopeptidase PaAP. While both prodrug motifs effectively neutralized the positive charge of D-Bac8C^Leu2,5^ and prevented DNA–peptide complex formation, EEEE□-Bac8C^Leu2,5^ showed negligible antimicrobial activity due to slow and incomplete activation. In contrast, ELEG□-Bac8C^Leu2,5^ underwent rapid PaAP□mediated activation, restoring bactericidal activity in planktonic cultures and biofilms. PaAP contributed significantly to complete prodrug activation, particularly within biofilms, where the accumulation of partially activated intermediates correlated with biphasic killing kinetics. The prodrug showed reduced activity against other ESKAPEE pathogens, demonstrating selective activation by *P. aeruginosa*. Experiments selecting resistant bacteria revealed distinct mutations in lipopolysaccharide biosynthesis pathways for D-Bac8C^Leu2,5^ and the prodrug, with limited cross□resistance. These findings establish aminopeptidase□activated AMP prodrugs as a promising approach for species□selective antimicrobial therapy and highlight the feasibility of exploiting bacterial enzymes for controlled antimicrobial peptide activation.

## Introduction

The threat of antimicrobial resistance (AMR) has been underestimated and is becoming a silent pandemic. In 2019, an estimated 6.22 million deaths were associated with AMR globally, and this is estimated to become 10 million by 2050.^1^ To highlight the urgency of this crisis, ESKAPEE bacteria (*Enterococcus faecium*, *Staphylococcus aureus*, *Klebsiella pneumoniae*, *Acinetobacter baumannii*, *Pseudomonas aeruginosa*, *Enterobacter species, and Escherichia coli*) were proposed by the World Health Organization (WHO) as priority targets for antimicrobial development. *Pseudomonas aeruginosa (P. aeruginosa)*, as an opportunistic and multidrug-resistant pathogen, can infect patients with cystic fibrosis, causing severe disease.^2,3^ Moreover, *P. aeruginosa* forms biofilms that contribute to resistance due to limited antibiotic penetration, reduced metabolic activity, and genotypic heterogeneity, enabling long-term survival during infection. Faced with the high rate of biofilm-related clinical infections and the lack of approved antibiofilm therapeutics, antimicrobial peptides (AMPs) show great promise as novel antibiofilm agents.^4^ However, instability and insufficient biosafety are barriers for new AMPs to enter clinical pipelines.^5^ A key objective in AMP design is to achieve high selectivity against pathogenic bacteria without exerting deleterious effects on host cells at therapeutic concentrations.^5^ While multiple AMPs have been designed to selectively inhibit bacterial protein targets and minimize host toxicity^6,7^, prodrug strategies offer an alternative approach to enhancing selectivity and reducing toxicity.^8,9^ Prodrugs are derivatives of therapeutic molecules that go through an enzymatic and/or chemical transformation *in vivo* to release the active parent drug, reestablishing the desired pharmacological effect.^10^

Previous prodrugs for cationic antimicrobial peptides consist of a positive charge-shielding moiety with negatively charged residues, a protease-specific cleavage site, and the AMP cargo. Upon exposure to activating enzymes, the shielding moiety of the AMP prodrug is removed to release the active peptide. As bacteria secrete proteases and peptidases for virulence and nutrient acquisition during infection, these proteins can be utilized as potential prodrug activation enzymes. While several AMP prodrugs have been designed to be activated by human neutrophil elastase (hNE)^11,12^ and human serum proteases^13,14^, few efforts have been made to exploit bacterial proteins to achieve AMP release upon exposure to bacteria^15^, particularly to ESKAPEE bacteria.^16^

**Scheme 1.**
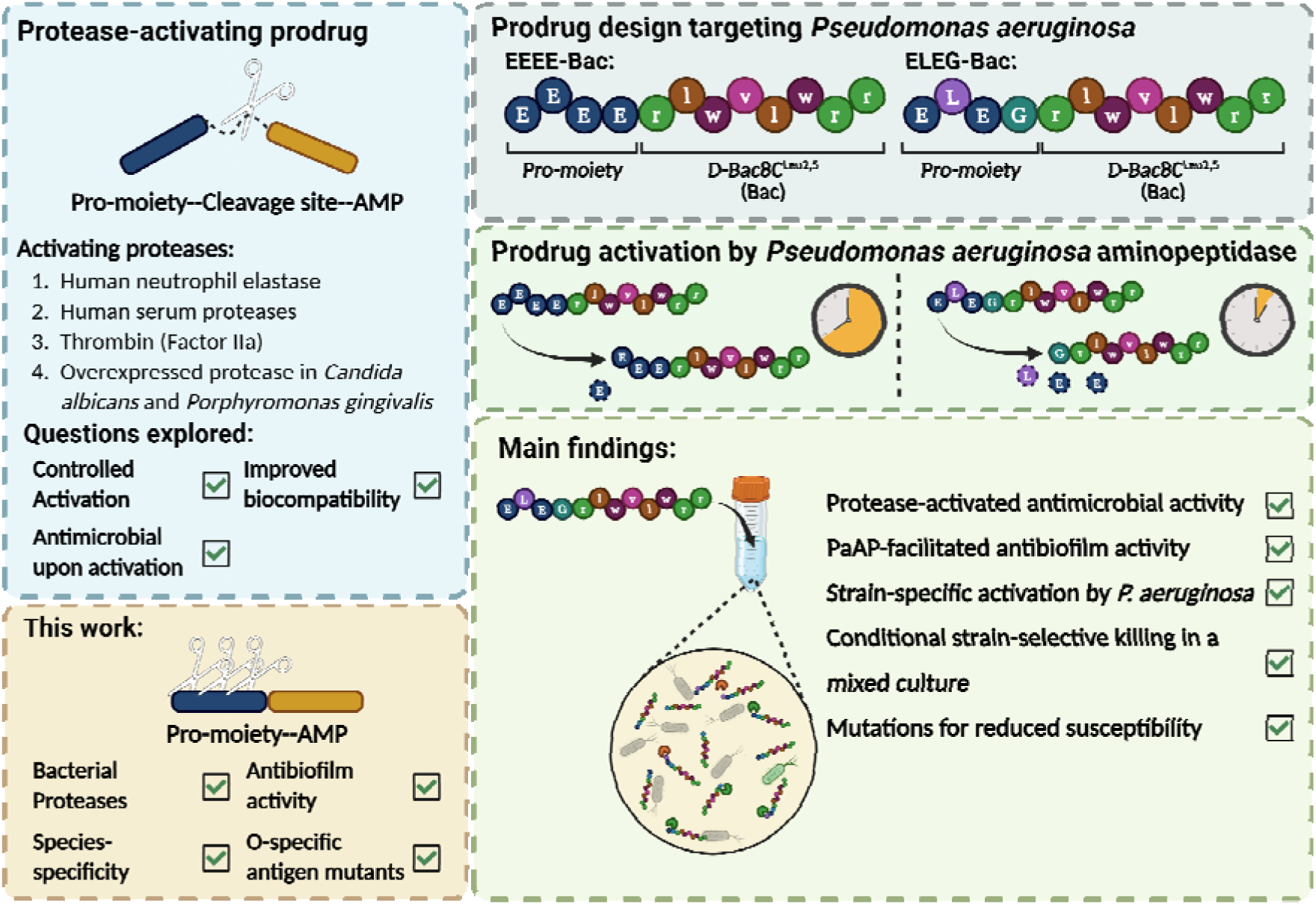
Design of bacterial aminopeptidase-activated antimicrobial peptide prodrug targeting *P. aeruginosa*.

To harness this bacterial proteolytic reservoir, we designed a prodrug moiety for a cationic antimicrobial peptide, D-Bac8C^Leu2,5^, targeting *P. aeruginosa* extracellular proteases as activating enzymes (Scheme 1 and Table 1). Bac8C^Leu2,5^ (RLWVLWRR-NH_2_), indicated as Bac^L^ below, was derived from bactenecin, the smallest naturally-occurring antimicrobial peptide identified in bovine neutrophils, and optimized by Hancock and colleagues in 2005.^17^ Unlike many cationic antimicrobial peptides that destabilize the outer membrane and induce cell lysis, Bac8C exerts bactericidal activity by depolarizing the cytoplasmic membrane potential, followed by inhibition of protein synthesis, eventually causing cell death without apparent cell lysis.^18,19^ D-Bac8C^Leu2,5^, indicated as Bac below, is composed of D-amino acids^20^, making it an ideal candidate for designing a prodrug AMP, given its protease insensitivity and broad-spectrum activity against several ESKAPEE bacteria.

**Table 1.** Antimicrobial peptides and prodrugs used in this study. L-amino acids and D-amino acids were written in upper-case letters and lower-case letters, respectively.

| Name | Sequence | Net charges |
| --- | --- | --- |
| Bac<sup>L</sup> | RLWVLWRR-CONH<sub>2</sub> | +4 |
| Bac | rlwvlwrr-CONH<sub>2</sub> | +4 |
| EEEE-Bac | EEEE-rlwvlwrr-CONH<sub>2</sub> | 0 |
| ELEG-Bac | ELEG-rlwvlwrr-CONH<sub>2</sub> | +2 |

## Methods

### 1. Media and strains

All chemicals were purchased from Fisher Scientific and Sigma-Aldrich. PaAP was expressed and purified in SHuffle® T7 Express Competent *E. coli* as previously described.^21^ Human neutrophil elastase (hNE) was purchased from Sigma-Aldrich. Lysogeny broth (LB) contained 10 g/L NaCl, 10 g/L tryptone, and 5 g/L yeast extract. For LB agar, LB was solidified with 15 g/L agar. Where needed in strain construction, carbenicillin was used at 100 ug/ml and gentamicin was used at 20 ug/ml. Jensen’s medium contained 85.6 mM NaCl, 14.4 mM K_2_HPO_4_, 92 mM L-glutamic acid, 24 mM L-valine, 8 mM L-phenylalanine, 70 mM glucose, 1.33 mM MgSO_4_, 0.14 mM CaCl_2_, 3.9 μM FeCl_3_, and 8.5 μM ZnCl_2_. Both LB and Jensen’s medium were filter sterilized before use.

*Pseudomonas aeruginosa* UCBPP-PA14, *Klebsiella pneumoniae* MGH 78578, and *Acinetobacter baumannii* ATCC19606 were obtained from Dr. Megan Bergkessel (University of Dundee). *Escherichia coli* ATCC 35218 was obtained from Dr. Jo Hobbs (University of St. Andrews). *Staphylococcus aureus* NCTC 8325 was purchased from the National Collection of Type Cultures (NCTC), UK Health Security Agency. The clean deletion of PaAP in the background of *Pseudomonas aeruginosa* UCBPP-PA14 strain (ΔPaAP) was carried out as previously described and used without further modification.^21^ WT UCBPP-PA14 marked with superfolder GFP (strain MB096) was previously described^22^. *E. coli* UTI89 marked with mScarlet was constructed as described below. The pUC18Tn7T Gm^R^ plasmid carrying the P_trc_ promoter^23^ was assembled with the CBR2 luciferase gene (sequence purchased as a gblock from IDT) and the mScarlet-I sequence from pMRE-Tn7-145 plasmid^24^using Gibson assembly (HiFi master mix, NEB) after amplifying the relevant sequences with the primers listed below (Q5 polymerase, NEB). From this plasmid, the cassette flanked by the Tn7R and Tn7L sequences was amplified by pMRE_movF and pMRE_movR primers and assembled by Gibson assembly into pMRE-Tn7-145 which had been digested with PacI and AvrII (NEB). This plasmid (stored as strain MB213) was electroporated into *E. coli* strain SM10 and then conjugated into *E. coli* UTI89 as previously described^24^ to yield strain MB215. Successful integration of the cassette and loss of the pMRE plasmid was verified by sensitivity to carbenicillin. Table S1 depicts all primers used.

For starting cultures, a single colony was used to inoculate 10 mL LB, and the cells were cultured at 37 ℃ while shaking at 180 rpm for 16-18 h.

### 2. Peptide synthesis

Antimicrobial peptide Bac (rlwvlwrr-NH_2_) and its prodrugs, ELEG-Bac and EEEE-Bac, were synthesized as previously described.^12^ All peptides were dissolved in water and stored at -20 ℃.

### 3. Electrophoretic mobility shift assay

Electrophoretic mobility shift assays were used to evaluate nonspecific DNA-AMP binding as previously described.^25^ Briefly, a dilution series of Bac, EEEE-Bac, and ELEG-Bac (0.5 μM to 64 μM) was added to 100 ng pJ411 plasmid (5157 bp) in 20 mM Tris, pH 7.5, and incubated at room temperature (RT) for 30 min before loading onto a 1% Tris-acetate-EDTA agarose gel pre-stained with SYBR Safe DNA gel stain (Thermo Fisher). Electrophoresis was carried out at 100 V for 60 min, and the gel was imaged using a Bio-Rad ChemiDoc^™^ MP imaging system. Similarly, the binding of a 326-bp DNA fragment to Bac, ELEG-Bac, Bac^L^, and EEEE-Bac was investigated using an 8% polyacrylamide gel made immediately before use. Sequences of the plasmid and the DNA fragments are provided in Table S2.

### 4. Enzymatic prodrug activation kinetics

For prodrug activation by PaAP, PaAP and prodrug peptides were added to the reaction buffer (50 mM HEPES, 0.2 M NaCl, pH 8.5) to final concentrations of 100 nM and 32 μM, respectively. For prodrug activation by hNE, 0.001 U hNE and 32 μM ELEG-Bac were added to PBS buffer (10 mM phosphate buffer, 2.7 mM KCl, 137 mM NaCl, pH 7.4). The reaction was incubated at 37 C.

Thirty microliter samples were collected from the reaction mixture and quenched with an equal volume of 2% trifluoroacetic acid (TFA). The mixture was centrifuged at 16000 x g for 15 min at RT to remove protein precipitate, and 60 μL of the supernatant was diluted with 60 μL of 0.1% formic acid. Caffeine, as an internal standard, was spiked into all samples to a final concentration of 10 μM before LC-MS analysis.

Ten microliters of each sample were injected into XSelect PREMIER^™^ HSS T3 (2.5 μm, 4.6 x 50 mm) column. To elute analytes, 0.1% formic acid in water (mobile phase A) and 0.1% formic acid in acetonitrile (mobile phase B) were used with a gradient (0-2 min: 1% B; 2-9 min: 1%-99% B in a linear gradient; 9-11 min: 99% B; 11-14 min: 1% B) at a flow rate of 0.4 mL/min. Ions of interest with selected *m/z* ratios were detected by a ACQUITY QDa detector (Waters) with distinct SIR (single ion reaction) channels for the parent peptide and each peptide intermediate for sequential cleavage events by PaAP. SIR peaks were integrated and peak areas were normalized by dividing each by the peak area of caffeine to correct for minor differences in injection volume. All m/z ratios and retention times for elution peaks used for quantification are listed in 3. All experiments were performed with two biological replicates and measured independently twice. Areas were averaged and plotted against time. The time-resolved curve was fitted with Equation 1 and 2.

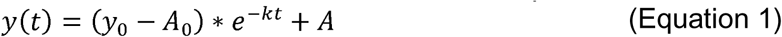

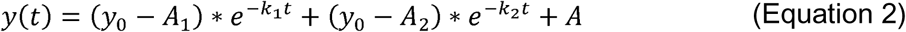

where y(t) is the normalized peak area at time t, y_x_-A_x_ is the amplitude of exponential phase for intermediate x, *k*_x_ is the rate constant of intermediate x, y_0_ is the value of y at time 0, A is the plateau after the exponential phase.

### 5. Leucine p-nitroanilide (Leu-pNA) assay

Cells from overnight cultures of tested strains were collected, washed with PBS, and diluted in 15 mL Jensen’s medium to OD of 0.05. Cultures were grown at 37 ℃ while shaking at 180 rpm. 1 mL of culture supernatant was collected by centrifugation at 16000 x g for 3 min, sterile filtered, and stored at -80 ℃ until further use. A leucine p-nitroanilide (Leu-pNA) stock solution of 200 mM was prepared in DMSO and further diluted in water to a final concentration of 4 mM. For aminopeptidase activity assays in supernatant, 50 μL of culture supernatant were added to 50 μL of 4 mM Leu-pNA in H_2_O with 2% DMSO, and the absorbance at 405 nm was monitored over 30 min at 25 C. All measurements were conducted with three technical replicates (independent experiments from one culture supernatant).

### 6. Minimum inhibitory concentration (MIC)

Cells from a 1 mL starting culture were pelleted by centrifugation at 8000 x g for 5 min, washed with 1 mL sterile PBS, and diluted in Jensen’s medium to OD of 0.05 before use. Antimicrobial peptides were added to cells in Jensen’s medium, and 200 μL were transferred to a sterile 96-well F-bottom plate (Greiner Bio-one). Cell growth was monitored by measuring OD_500_ in a Tecan plate reader at 37 ℃ while shaking at 240 rpm for 24-48 h. All experiments were performed with three biological replicates. The MIC for the prodrug in this study refers to the minimum prodrug concentration at which there is no increase in optical density after the initial growth phase.

### 7. Minimum biofilm eradiation concentration (MBEC_90_)

To establish mature biofilms, 50 μL of cell suspension of WT and ΔPaAP strain at an OD_600_ of 0.05 in Jensen’s medium were added to a sterile 96-well flat-bottom plate and incubated at 37 C. After 24 h, all culture supernatants were removed, and biofilms were washed with 100 μL of sterile PBS twice. A series of Bac and ELEG-Bac in Jensen’s medium were added to the biofilm, and the plate was incubated at 37℃ for a further 24 h.

After washing each well twice with 100 μL of sterile PBS, the total metabolic activity was measured by adding 60 μL of BacTiter-Glo™ Microbial Cell Viability Assay reagent (Promega) diluted in sterile PBS to the biofilms (1:1, *v/v*). The plate was incubated at RT for 5 min to lyse cells, and 50 μL was transferred to a white 96-well half-area plate (Greiner Bio-one). The bioluminescence intensity was measured in a Tecan infinite 200 Pro plate reader. All experiments were conducted with four biological replicates. EC50 value was determined with a dose-dependent sigmoidal model (Equation 3).

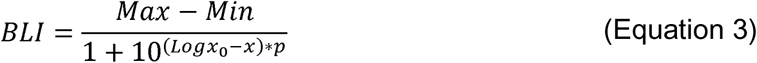

### 8. Propidium iodide (PI) staining of biofilms

Mature biofilms were grown according to the method described below. Culture supernatant from 1-day-old biofilms was removed, and biofilms were washed twice with 100 μL of sterile PBS. A 3 mM PI stock was prepared in DMSO, and the stock was diluted in Jensen’s medium to reach the final concentration of 30 μM, and 59.4 μL were added to each well. The plate was incubated at 37 C for 15 min. 0.6 μL of a 100x Bac or ELEG-Bac stock was added to biofilms, and PI fluorescence was monitored at 37 C for 10 h in a Tecan plate reader using an excitation filter at 485 ± 20 nm and an emission filter at 620 ± 10 nm. Curves from Bac-treated biofilms were fitted with a single exponential equation. A biphasic curve (Equation 4) was used to fit the fluorescence intensity from prodrug-treated biofilms of the WT strain. All experiments were conducted with three biological replicates.

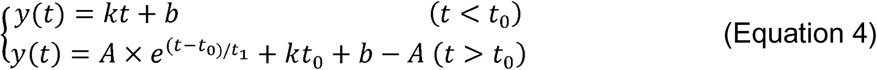

where t_0_ is the transition points of the biphasic curve, t_1_ is the time constant of the exponential phase, k is the slope of the linear phase, A is the amplitude of the exponential phase.

### 9. Identification of prodrug activation intermediates

ELEG-Bac was added to the cell culture of WT and ΔPaAP strain in Jensen’s medium to the final concentration of 32 μM, and the culture was incubated at 37 C with shaking. After 24 h, the cell culture was collected and heated at 95 ℃ for 5 min to kill all bacteria and denature most proteins, followed by acidification using an equal volume of 2% TFA to quench further activation. The sample was vortexed and centrifuged at 16000 x g for 15 min.

Samples were subjected to LC-MS/MS using an Ultimate 3000 RSLC (Thermo Scientific) coupled to an Orbitrap Fusion Lumos mass spectrometer (Thermo Scientific). Samples were injected onto a reverse-phase trap (Pepmap100 C18 5μm,100A, 100um × 2cm) for pre-concentration and desalted with loading buffer (100% water, 0.05% TFA), at 5 μL/min for 10 min. The peptide trap was then switched into line with the analytical column (Easy-spray Pepmap RSLC C18 2um, 15cm x75um ID). Peptides were eluted from the column using a linear solvent gradient (Mobile phase A: 100% water, 0.1% formic acid, Mobile phase B: 80% acetonitrile, 20% water, 0.1% formic acid) using the following gradient: linear 2–35% of buffer B over 50 min, sharp increase to 95% buffer B within 0.5min, isocratic 95% of buffer B for 5 min, sharp decrease to 2% buffer B within 1min and isocratic 2% buffer B for 5 min. The mass spectrometer was operated in DDA positive ion mode with a cycle time of 1.5 sec. The Orbitrap was selected as the MS1 detector at a resolution of 60000 with a scan range of from m/z 375 to 1500. Peptides with charge states 2^+^ to 5^+^ were selected for fragmentation in the Ion trap using HCD at 35% normalized collision energy.

### 10. Cell viability assay

Cell viability was measured according to a previously reported protocol.^26^ Briefly, 100 μL 3 × 10^5^ cells/mL HEK293FT cells were seeded in a 96-well flat-bottom transparent plate and was incubated at 37 C for 24 h. Cells were washed with fresh 100 μL DMEM twice. A series of 100x stock solutions of Bac and ELEG-Bac was prepared in DMSO and added to a final 1x concentration. 1% Triton X-100 was used as a positive control. The plate was incubated for 24 h before staining with 1.2 mM Thiazolyl Blue tetrazolium bromide in DMEM for 2 h at 37 C. Insoluble MTT formazan was solubilized by adding 100 μL isopropanol acidified with 1.5 M HCl and aspirating with a pipette. Absorbance at 540 nm was measured in a Tecan plate reader. Relative viability was calculated using Equation 5.

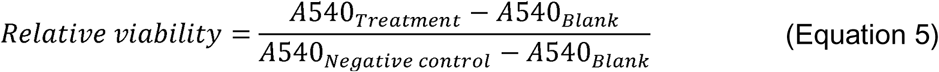

### 11. Antimicrobial test of the prodrug in a coculture of GFP-labelled *P. aeruginosa*

WT strain and mScarlet-labelled *E. coli* UTI89 strain 1 mL overnight culture of both strains were diluted to an OD_600_ of 0.1 in Jensen’s medium supplied with 10% LB (*v/v*). The culture was mixed at different ratios to reach the proportion desired of both strains while keeping the same cell density. ELEG-Bac was added to 200 μL mixed culture in a 96-well μclear^®^ plate (black with transparent bottom). The plate was incubated at 37 C for 7 minutes to reduce condensation. The optical density and fluorescence were monitored in a Tecan plate reader for GFP fluorescence (excitation filter: 485 ± 20 nm; emission filter: 535 ± 25 nm) and for mScarlet fluorescence (excitation filter: 560 ± 20 nm; emission filter: 620 ± 10 nm).

### 12. Identification and quantification of GFP-labelled *P. aeruginosa* and mScarlet-labelled *E. coli* from the mixed culture

Mixed cultures at different mixing ratios of *P. aeruginosa* and *E. coli* were serially diluted and inoculated on LB agar dishes at the desired cell density. The plate was incubated at 37 C for 24 hours and was imaged using a LICOR Odyssey^®^ M Imaging System using 488 channel (green; excitation: 488 nm; emission: 519-543 nm) and 520 channel (red; excitation: 520 nm; emission: 570-610 nm). Images were adjusted to reduce background noise. Images at green channel, red channel, and the merged channel were exported in tiff format.

CFU was counted manually, and images at the merge channel were analyzed using OpenCFU for GFP-labelled *P. aeruginosa* WT strain.

### 13. Selection of strains with reduced susceptibility and whole genome sequencing

Strains were exposed to an increasing AMP concentration (0.5x MIC, 0.75x MIC then to 1x MIC). Briefly, Bac and ELEG-Bac were added to the cell dilution in Jensen’s medium to the final concentration of 0.5x MIC. OD_500_ was monitored at 37 C while shaking at 360 rpm for 24 h. Replicates with visible growth were diluted into fresh medium containing 0.75x MIC AMP or the prodrug, and subsequently, 1x MIC at the ratio of 1:50.

At 1x MIC, all replicates with visible growth were stored as glycerol stocks and used to inoculate LB agar plates.

To confirm the reduced antimicrobial susceptibility, 6 colonies were randomly selected from LB agar plates and used to inoculate 10 mL LB. Cells were collected by centrifugation at 8000 x g for 5 min and diluted in Jensen’s medium to an OD_600_ of 0.05. Bac or ELEG-Bac was added to 1x MIC, and cells were grown for two more passages. Thereafter, cells were diluted in 10 mL Jensen’s medium and were cultured at 37 C for 8 h while shaking at 180 rpm. Cells were then collected by centrifugation at 4000 rpm for 20 min and were resuspended in 500 μL DNA/RNA shield (Zymo Research) for Illumina Long Reads performed by MicrobesNG (United Kingdom) with target coverage of 50x . Raw Illumina reads in FASTQ format were assembled and analyzed with breseq^27^, using *Pseudomonas aeruginosa* UCBPP-PA14 complete genome from GenBank as the reference genome.

## Results

### Converting cationic AMP into prodrugs decreases the overall positive charge and prevents formation of DNA-AMP complexes

As DNA is negatively charged under physiological conditions, it can form complexes with cationic antimicrobial peptides, which can be visualized by DNA band shifts on agarose gels.^28,29^ These complexes often reduce the potency of cationic AMPs, effectively quenching their biological activity. To test if the prodrug moiety alters the electrostatic properties of Bac, an electrophoretic mobility shift assay (EMSA) was used to probe its electrostatic state by evaluating the DNA binding capability of Bac and its associated prodrugs.

DNA band shifts were observed with Bac at concentration from 8 to 64 μM (Figure 1A), suggesting the formation of Bac-plasmid complex with retarded mobility. To test the specificity, Bac and Bac^L^ were used to interact with distinct DNA sources, and we observed DNA band shifts in all four combinations, indicating non-specific interactions (Figure 1B and Figure S1). To investigate the major stabilizing force of the complex, we increased the ionic strength of the buffer, and this abolished the DNA band shift (Figure 1C), demonstrating that electrostatic interactions play a major role in stabilizing the AMP-DNA complex.

**Figure 1.**
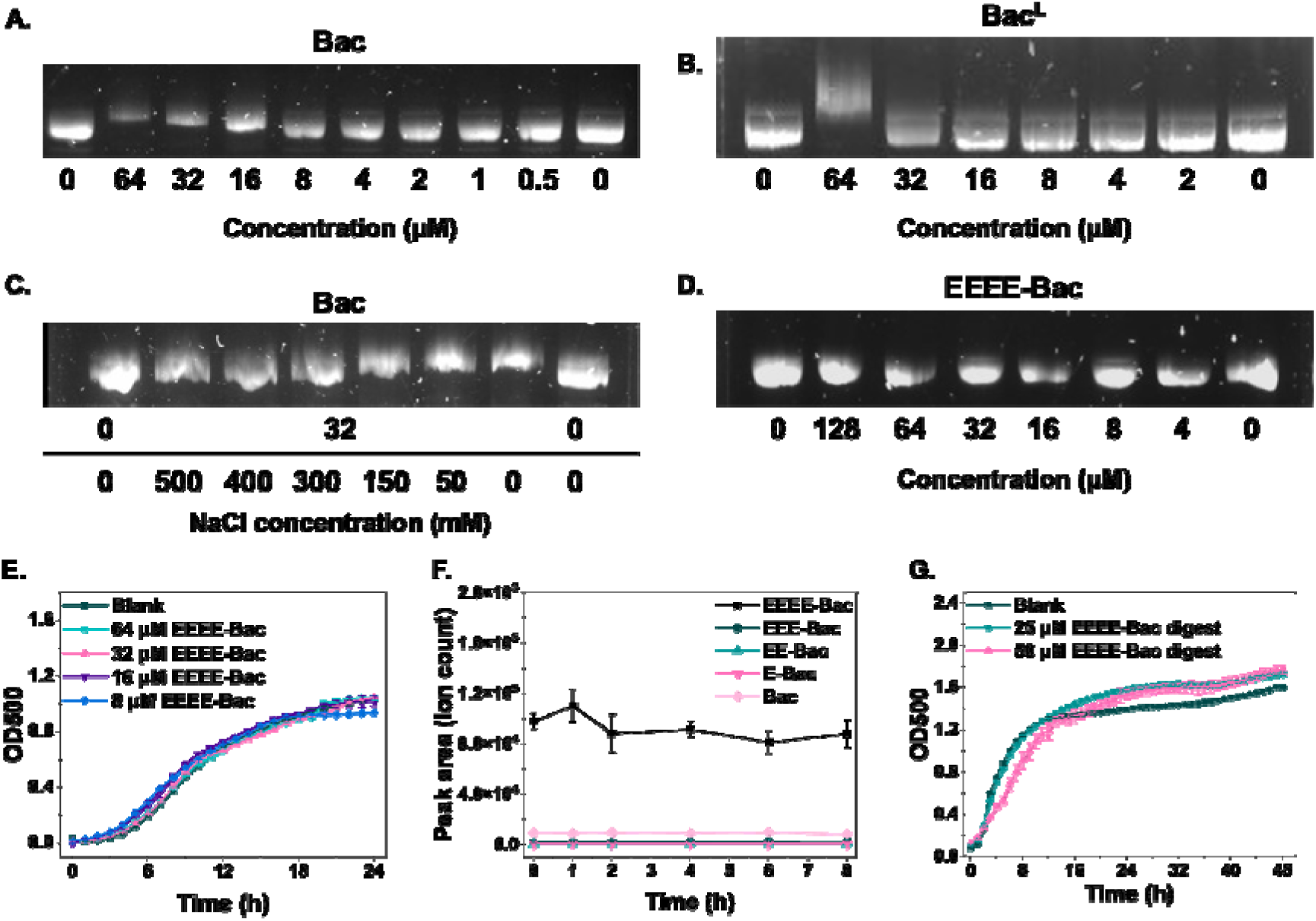
The prodrug EEEE-Bac neutralizes the positive charge and inactivates Bac but does not inhibit the growth of *P. aeruginosa*. (A-D) DNA agarose gel imaging revealing the interaction between a pJ411 plasmid and (A) Bac; (B) Bac^L^; (C) 32 μM Bac in the presence of NaCl; and (D) EEEE-Bac. Full gels are depicted in Figure S1; (E) growth curve of *P. aeruginosa* PA14 strain in the presence of EEEE-Bac; (F) the activation kinetics of 32 μM EEEE-Bac by 100 nM PaAP at 37 ℃; (G) growth curve of *P. aeruginosa* PA14 strain in the presence of three-day digestion of 50 μM EEEE-Bac by PaAP.

To convert Bac into a prodrug, an N-terminal sequence with four L-glutamic acids was added to the parent AMP to reduce the net positive charge from a calculated +4 to 0. As proof of concept, we evaluated the formation of prodrug-DNA complex by EMSA, and no DNA band shift was observed with up to 128 μM EEEE-Bac (Figure 1D), establishing that the prodrug sequence can shield positive charges of Bac.

### EEEE-Bac cannot be fully activated by *P. aeruginosa* and does not show anti-*P. aeruginosa* activity

The antimicrobial activity of EEEE-Bac was tested in *Pseudomonas aeruginosa* UCBPP-PA14 (PA14) strain, and the prodrug did not show antimicrobial activity against *P. aeruginosa* over 24 hours (Figure 1E). *in silico* cleavage site predication indicated that staphylococcal peptidase I (SspA), Asp-N endopeptidase, and proteinase K may be able to cleave the prodrug sequence. While two proteins, PA14_47090 and PA14_18630, were identified as homologues of SspA and proteinase K in *P. aeruginosa* PA14 proteome, respectively, the sequence similarity of their peptidase domains to the two reference proteases is below 30%, and neither was predicted to be extracellular proteases by PSORTb 3.0 (Figure S2). *P. aeruginosa* secretes the peptidase Mep72 with Asp-N endopeptidase activity; however, its activity is biofilm-specific, and it is thought to process a restricted set of virulence factors exported by the type III secretion system, and therefore it may not contribute to the prodrug activation in planktonic cultures.^30,31^

Our recent work on the extracellular aminopeptidase from *P. aeruginosa* aminopeptidase (PaAP) showed that it can remove amino acids from the N-termini of unstructured peptides.^21^ Although PaAP is not constitutively expressed in *P. aeruginosa*, the gene (*lap*) is found in 7,976 of 7,982 *Pseudomonas aeruginosa* genomes deposited in the Pseudomonas Genome Database (https://pseudomonas.com/), and its activity is detectable in Jensen’s medium after 8 hours of growth (Figure S3). The expression of PaAP is under the control of SutA and RpoS^22,32^, two transcriptional regulators to response to stress. PaAP is one of the most abundant peptidases in the extracellular proteome in the planktonic culture in a minimal glucose medium and colony biofilm matrix on LB agar.^33,34^. Moreover, PaAP contributes to virulence by facilitating biofilm formation in a *P. aeruginosa*-lung epithelial cell co-culture model^35^, and it has been identified in sputum from cystic fibrosis patients^36^, highlighting its relevance during infection and long-term colonization.

Using purified PaAP to test the activation kinetics of EEEE-Bac showed no significant activation over an eight-hour incubation at 37 C (Figure 1F). Within 72 h of incubation, approximately 50% EEEE-Bac was processed. This indicates very slow activation and incomplete removal of the prodrug sequence by PaAP (Figure S4). Furthermore, this PaAP digest product did not exhibit bactericidal effects in *P. aeruginosa* PA14 culture over 48 hours (Figure 1G).

### ELEG-Bac shows antimicrobial activity against planktonic and biofilm cultures of *P. aeruginosa*

Previous protease-activated prodrug designs utilize multiple aspartic acids and glutamic acids to shield and neutralize positive charges. However, this strategy was incompatible with our design, where ideally the prodrug would be activated by multiple *P. aeruginosa* PA14 proteases in the absence of a protease-specific cleavage site ( 1). Thus, a better prodrug sequence was required to accelerate activation and avoid potential activity-quenching effects of the remaining L-glutamic acid at the N-terminus of Bac.

Based on prior work on the substrate scope of PaAP, we designed the improved prodrug ELEG-Bac with two positive net charges with reduced binding affinity to DNA (Figure S1). We hypothesized that this prodrug sequence would be efficiently cleaved by PaAP, leaving a “glycine scar” before the AMP sequence.

*In vitro* activation of 32 μM ELEG-Bac with 0.1 μM PaAP revealed a half-life of 23 minutes (Figure 2A), and the final product of the activation was G-Bac.

**Figure 2.**
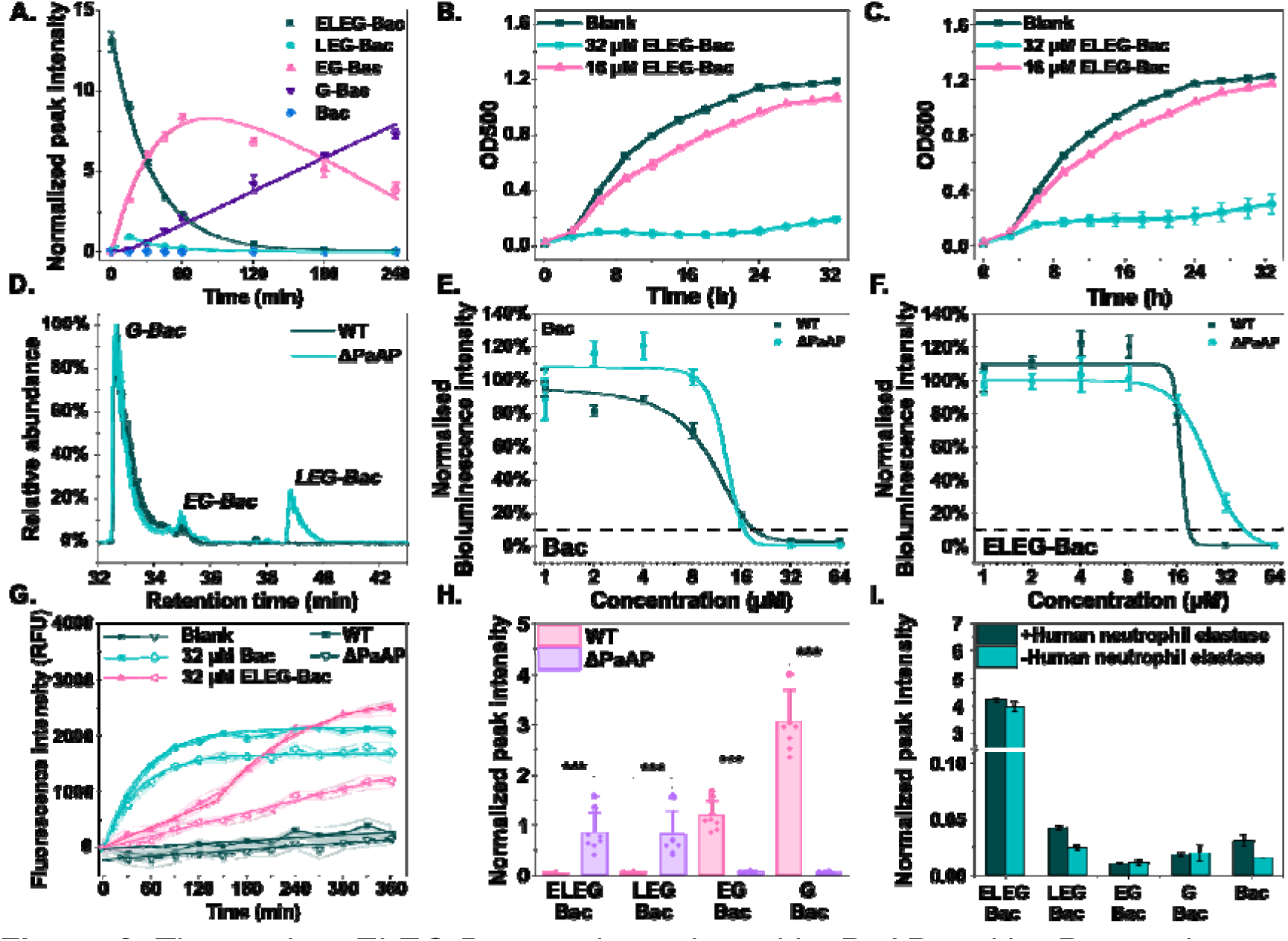
The prodrug ELEG-Bac can be activated by PaAP and by *P. aeruginosa* PA14 strain *in vitro* and shows antimicrobial activity in planktonic culture and biofilms. (A) Relative quantification of intermediates in the prodrug activation experiment with PaAP. Data points were fitted with Equation 1 and Equation 2, and fitted parameters were listed in Table S4; (B & C) growth curves of (B) WT and (C) ΔPaAP strain in the presence of the prodrug; (D) identification of prodrug activation intermediates in the culture of WT and ΔPaAP after 24 h exposure to ELEG-Bac by tandem mass spectrometry. Chromatograms including the elution peaks of prodrug intermediates were shown with labelling for each elution peak; (E & F) metabolic activity of WT and ΔPaAP biofilms after 24 h treatment of (E) Bac and (F) the prodrug. Data were normalized against the average bioluminescence intensity without Bac or ELEG-Bac treatment. Dashed line indicates 10% metabolic activity as the threshold of MBEC_90_. Data were fitted with a sigmoidal equation to estimate the breakpoint.; (G) propidium iodide staining of WT and ΔPaAP biofilms in the presence of 32 μM Bac or 32 μM ELEG-Bac at 37 ℃. The biphasic curve was fitted with Equation 4, and the fitted parameters are listed in Table S5. The blank refers to the growth in Jensen’s medium without ELEG-Bac; (H) quantification of intermediates in the supernatant from WT and ΔPaAP biofilms after 24 h treatment of 32 μM ELEG-Bac at 37 (*: p<0.05; **: p<0.01; ***: p<0.001); ℃ (I) digestion of 32 μM ELEG-Bac by 0.001U human elastase at 37 ℃ in 10 mM PBS saline (pH 7.4).

Given the rapid activation kinetics mediated by PaAP, we tested the antimicrobial activity of ELEG-Bac against *P. aeruginosa* (Table 2). From growth curves, an inhibition lasting 16 hours was followed by slow recovery observed after prolonged growth times. This indicated that the prodrug can be activated and inhibit the growth of *P. aeruginosa* (Figure 2B).

**Table 2.** Minimum inhibitory concentration of Bac and ELEG-Bac in the planktonic culture of tested ESKAPEE bacteria.

| *P. aeruginosa* UCBPP-PA14 | Bac<br>(μM) | ELEG-Bac<sup>a</sup><br>(μM) |
| --- | --- | --- |
| WT | 16 | 32 |
| ΔPaAP | 16 | 32 |
| **Other Gram-negative** |  |  |
| *A. baumannii* ATCC 19606 | 16 | >64 |
| *K. pneumoniae* MGH 78578 | 16 | >64 |
| *E. coli* ATCC 25922 | 8 | 64<sup>b</sup> |
| **Gram-positive** |  |  |
| *S. aureus* NCTC 8325<sup>c</sup> | 8 | >64 |

<sup>a</sup> MIC for the prodrug is defined as the lowest peptide concentration causing no OD increase after the initial growth.
<sup>b</sup> There is no growth prior to the growth inhibition.
<sup>c</sup> LB was used in the test.

To evaluate the contribution of PaAP to the activation in the presence of bacteria, we monitored the growth of ΔPaAP strain with 32 μM ELEG-Bac, and the prodrug still led to growth inhibition against the ΔPaAP strain. There were no changes in MIC values, but a 2-hour delay in the growth inhibition compared to the WT strain (Figure 2C) as well as the accumulation of LEG-Bac after 24 h growth were observed in ΔPaAP cultures (Figure 2D and Figure S7). Since LEG-Bac did not accumulate in the PaAP activation experiment (Figure 2A), we propose that PaAP is crucial to fully remove the pro moiety and lead to G-Bac.

Additionally, the antibiofilm activities of Bac and the prodrug were investigated. While Bac showed a similar MBEC_90_ and EC50 in the WT biofilms (10.4 ± 1.4 μM) and ΔPaAP biofilms (12.7 ± 1.3 μM) (Figure 2F), we observed an elevated MBEC_90_ (64 μM) and EC50 (25.9 ± 0.6 μM) for ELEG-Bac in the ΔPaAP biofilms compared to WT biofilms (MBEC_90_ of 32 μM and EC50 of 16.4 ± 0.02 μM, Figure 2G). We hypothesize that the prodrug may not be completely activated in ΔPaAP biofilms and generate intermediates that are less lethal to biofilm cells. We used propidium iodide (PI) to track dead cells, as when cell membranes are permeabilized, PI can diffuse into the cytoplasm and bind to DNA. PI staining revealed an exponential increase of PI fluorescence upon the exposure of 2x MIC

Bac (Figure 2G). When treating WT biofilms with 32 μM ELEG-Bac, we observed a biphasic PI fluorescence curve, consisting of a slow linear increase followed by a rapid exponential increase for a fast-killing phase. In contrast, the fast exponential killing phase was not observed in ΔPaAP biofilms over 6 h. When quantifying the amount of prodrug intermediates in the supernatant of biofilm cultures, we found that EG-Bac and G-Bac were significantly less abundant in ΔPaAP biofilms in comparison to WT biofilms (Figure 2H). We propose that distinct intermediates contribute to each killing phase, with ELEG-Bac and LEG-Bac contributing to the slow-killing phase and EG-Bac and G-Bac contributing to the fast-killing phase.

Although the prodrug was resistant to cleavage by human neutrophil elastase (Figure 2I), the pro-moiety is not stable in human serum with a half-life of 40 minutes (Figure S5A). Comparing with the high stability in plasma (Figure S4B), we hypothesized that the non-specific activation in human serum could be attributed to metalloproteases in serum, such as human matrix metalloproteases (MMP) with similar substrate scopes with the most abundant proteases of *P. aeruginosa*, LasB.^37^ We did not observe a significant difference in cell viability of HEK293 FT cell lines after treatment with Bac or the prodrug for 24 hours (Figure S6). This may be attributed to the low cytotoxicity of Bac.^38^

### Prodrug AMP converts Bac from a broad-spectrum AMP into a narrow-spectrum antimicrobial compound

As prodrug activation depends on extracellular proteolytic activities, especially aminopeptidases, we investigated the antimicrobial properties of ELEG-Bac against four strains of ESKAPEE bacteria (Table 2 and Figure S8). ELEG-Bac showed low antimicrobial activity against *K. pneumoniae*, *E.* c*oli*, and *S. aureus*. The absence of prodrug intermediates in the culture supernatant suggested that *K. pneumoniae* and *E. coli* cannot activate the prodrug, whereas *S. aureus* can remove the glutamic acid from the N-terminus to generate LEG-Bac. This is likely due to the glutamyl endopeptidase activity of SspA (Figure 3A). A decrease of 20% in metabolic activity and increased PI fluorescence was observed in the culture of *A. baumannii* with 32 μM ELEG-Bac (Figure S9A-C), but compared to *P. aeruginosa*, the prodrug showed limited activation in *A. baumannii* (Figure S9D).

**Figure 3.**
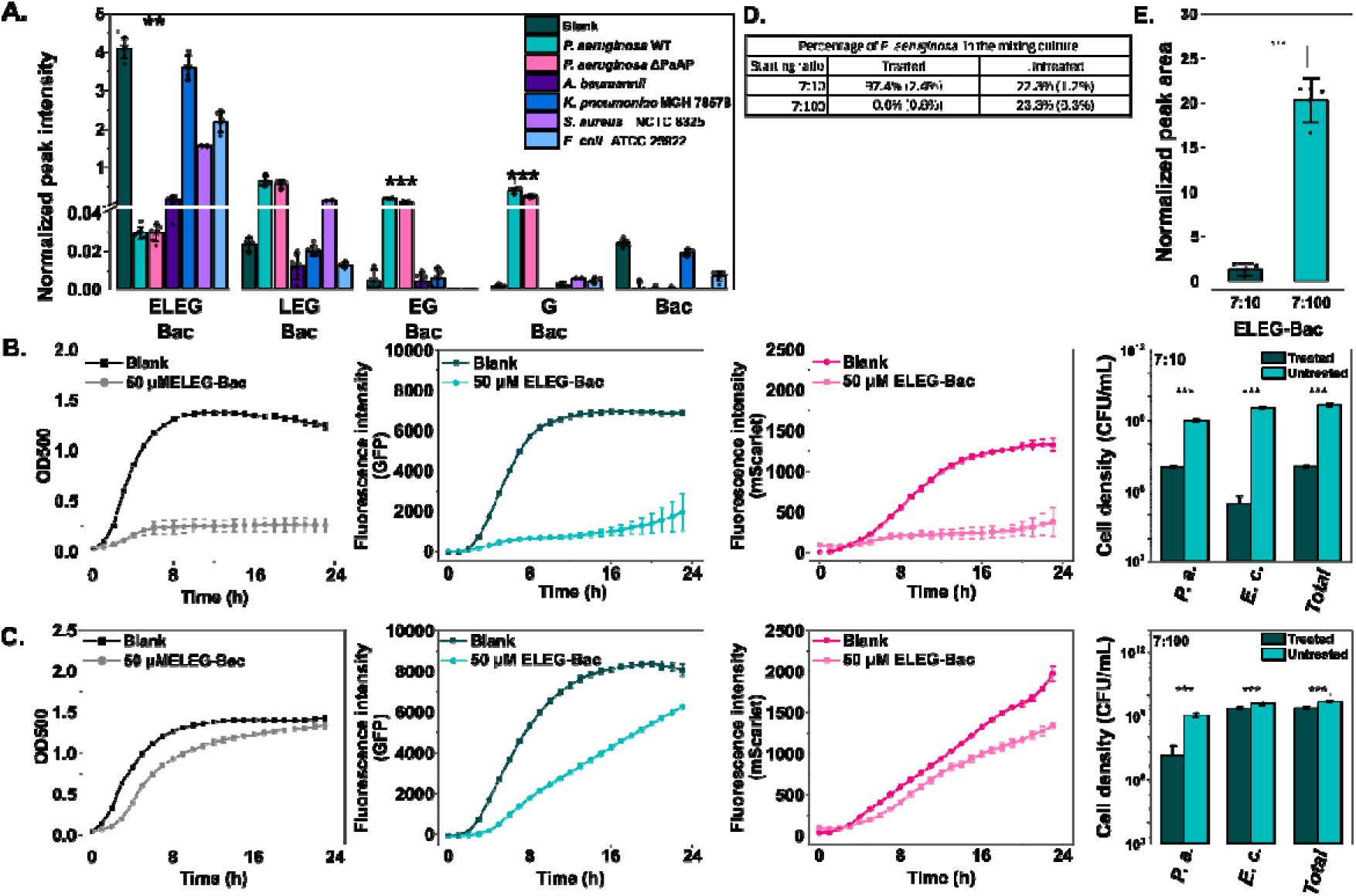
Prodrug is efficiently activated by *P.* aeruginosa, and not by other ESKAPEE bacteria. (A) Relative quantification of intermediates in the bacterial culture supernatant of different bacteria after 24-hour treatment with 32 μM ELEG-Bac. Blank is ELEG-Bac in Jensen’s medium without any inoculum; (B-C) growth of GFP-labelled *P. aeruginosa* WT and mScarlet-labelled *E. coli* UTI89 strains with 50 μM ELEG-Bac at the mixing ratio of (B) 7:10 and (C) 7:90 (cell count). The blank control refers to growth without addition of ELEG-Bac. Panels from left to right are growth curves in optical density (black), fluorescence from GFP-labelled *P. aeruginosa* WT strain (teal), mScarlet-labelled *E. coli* UTI89 strain (pink), and colony forming units of *P. aeruginosa* (P. a.), *E. coli* (E. c.), and total cells (Total) at the end of the experiment (bar graphs). Representative LB agar images in green channel, red channel, and merged images for quantification were shown in Figure S12; (D) relative quantification of the prodrug after the experiment in panel B and panel C (*: p<0.05; **: p<0.01; ***: p<0.001). All experiments were carried out as triplicates and data are shown as average and standard deviation.

Polymicrobial infections are common in catheter-associated urinary tract infections, where *P. aeruginosa* and *E. coli* can coexist.^39^ Considering the broad-spectrum antimicrobial activity after activating the prodrug, we questioned if the prodrug can be activated by *P. aeruginosa,* causing antimicrobial effects to *E. coli* when both grow in coculture. Growth of both bacterial populations remained stable in a coculture of *P. aeruginosa* PA14 and an *E. coli* UTI89 strain at a ratio from 7:10 to 7:100 (P*. aeruginosa* PA14:*E. coli* UTI89, Figure S10 and S11 show ratio of *P. aeruginosa* and *E. coli* in the mixed culture).

With 50 μM ELEG-Bac, growth arrest of both strains was observed at the mixing ratio of 7:10, which was absent when the mixing ratio to 7:100 was evaluated (Figure 3B-C). The prodrug was not activated when *E. coli* predominated in the coculture (Figure 3D), but successful cross-activation occurred at the 7:10 ratio.

### Mutations affecting lipopolysaccharide biosynthesis reduced susceptibility of *P. aeruginosa* to Bac and ELEG-Bac

After 24h growth in the presence of Bac or the prodrug, the wild type of *P. aeruginosa* restarted growing (Figure 2B and Figure S8A), suggesting a reduced susceptibility of Bac and ELEG-Bac. To investigate mutations leading to the reduced susceptibility, we increased the selective pressure by increasing Bac and ELEG-Bac concentrations from 0.5x MIC to 1x MIC. Cells were inoculated on LB agar plates. To explore whether mutations are shared by both strains, six colonies were randomly picked up for antimicrobial susceptibility test with 1x MIC Bac or ELEG-Bac in Jensen’s medium, followed by diluting and regrowing them for another two passages in the presence of 1x MIC Bac or the prodrug before sequencing (Figure 4A). In this way, we harvested six strains with reduced susceptibility to Bac (AR1-AR6), where the MIC was elevated to 32 μM (Figure S13A), and four prodrug strains with reduced susceptibility to the prodrug (PR1, PR2, PR5, and PR6).

**Figure 4.**
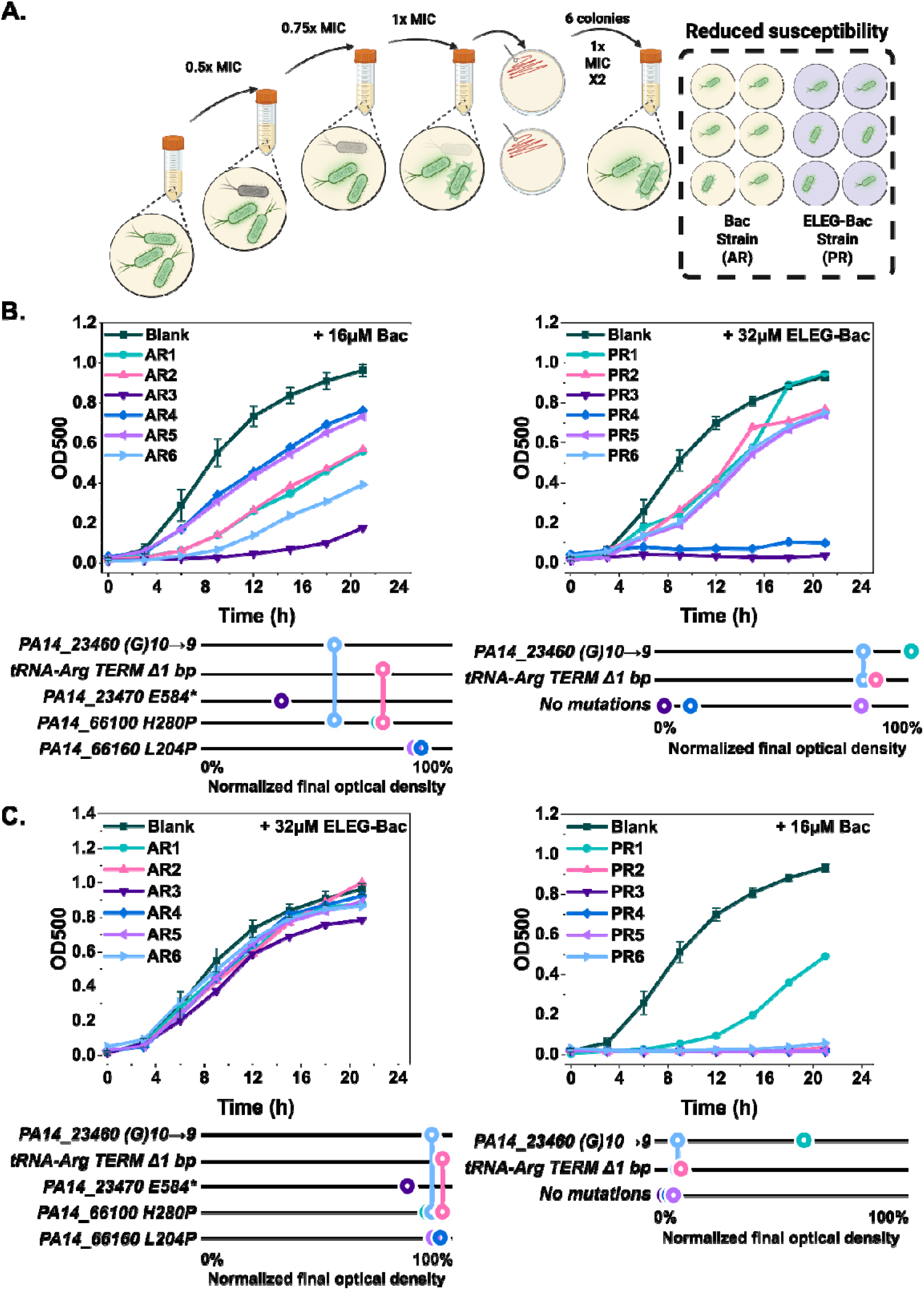
*P. aeruginosa* PA14 showed reduced susceptibility to Bac and ELEG-Bac with mutations in genes encoding four distinct glycosyltransferases. (A) Scheme for selecting strains with reduced susceptibility of Bac (AR1-AR6) and the prodrug (PR1-PR6) by increasing selective pressure; (B) growth curves of six strains treated with 16 μM Bac (left) and 32 μM ELEG-Bac (right); (C) cross-resistance experiments, where AR strains were grown in the presence of 32 μM prodrug (left) and PR strains were grown in the presence of 16 μM Bac (right). Identified gene mutations are shown under each graph, with the scale demonstrating the relationship between mutations and the final optical density of the culture compared to the wild-type blank. Each circle on the scale depicts the corresponding strain in the growth curve. Blank in panel B and C refers to the growth of WT strain without Bac or ELEG-Bac treatment.

Genome sequencing revealed five gene mutations related to the reduced susceptibility to Bac, and two shared mutations were also observed in PR strains. (Table 3) Except Rho-independent tRNA-arginine transcriptional terminator, all four proteins are related to the integrity of lipopolysaccharide (LPS). These mutations did not result in growth impairment in the absence of Bac or the prodrug (Figure S13 B).

**Table 3.**
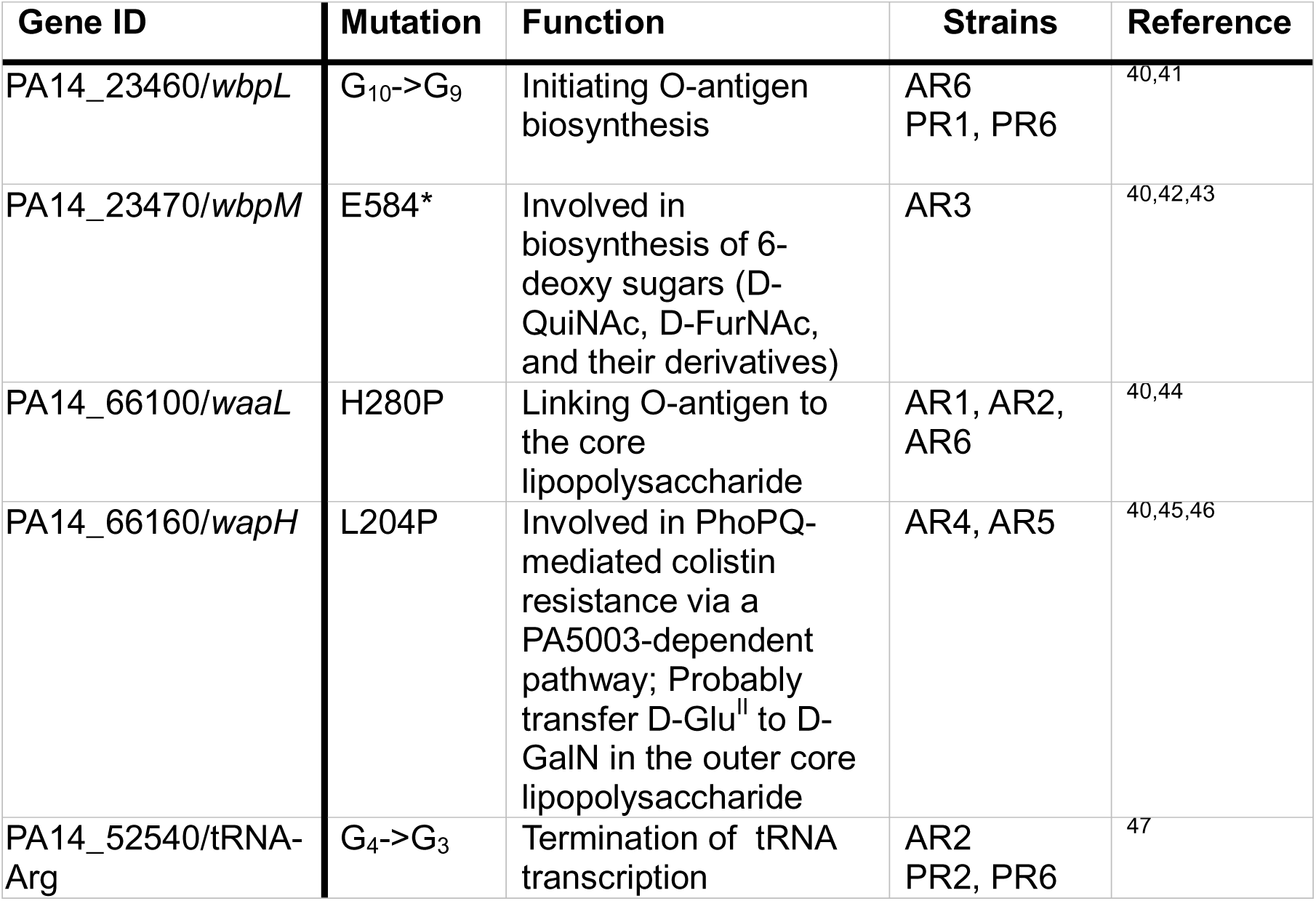
Identified mutations in AMP and prodrug strains.

A base pair loss was identified at the Rho-independent transcriptional terminator of tRNA-Arginine with an anticodon ACG (PA14_52540). Although this mutation supported bacteria to adapt to AMP and prodrug treatments, it does not add to the reduced susceptibility when other mutations coexist in the mutant. The transcription factor is an intrinsic terminator that can develop a hairpin structure and destabilize the transcriptional complex, terminating transcription. The loss of one guanine could extend the RNA hairpin structure from 9 base pairs to 12 base pairs, which significantly decreases the termination efficiency both in vitro and *in vivo*.^48^ As no mutations were identified at the other copy of tRNA-Arginine, it is unclear how this mutation contributes to the survival of PR strains.

Additionally, two AR strains with the highest reduction of AMP susceptivity acquired a single point mutation of ORF PA14_66160 (*wapH* in PAO1 strain) encoding a glycosyltransferase for core oligosaccharide biosynthesis.^40^ Another two strains shared a single amino-acid mutation of ORF PA14_66100 (*waaL* in PAO1 strain) that ligates O-antigen to the core polysaccharide of LPS.^44^ Sequence alignment of PA14_66100 homologues from other species revealed that mutations were observed at H280 (Figure S14A), while L204 in PA14_66160 is highly conserved across all sequences (Figure S14B). In addition to the same mutation in PA14_66100, AR6 lost one base pair at the coding region of ORF PA14_23460 (*wbpL* in PAO1 strain) encoding a glycosyltransferase that initiates biosynthesis of O-specific antigen in PA14 strain. The loss of guanine leads to a frameshift, changing the amino acid sequence from position 49 and truncating protein translation at position 53. Another strain with almost 12h growth inhibition acquired a mutation in ORF PA14_23470 encoding a conserved D-GlcNAc 4,6-dehydratase for short O-antigen biosynthesis across 20 *P. aeruginosa* serotypes.^40^

For PR variants, two strains did not show reduced susceptibility, and no mutations were identified in these two strains (Figure 4B). The nonsense mutation on ORF PA14_23460 was identified in two PR strains, suggesting that this mutation may be crucial for *P. aeruginosa* to reduce susceptibility of Bac and ELEG-Bac. This mutation is related to reduced susceptibility to beta-lactam antibiotics, and these may due to elevated hydrophobicity of the cell membrane, which eventually facilitates cell aggregation.^41,49^

Next, we tested if both strains also showed reduced susceptibility to the other compound in a cross-resistance experiment. All AR strains showed no growth impact in the presence of prodrug (Figure 4C), suggesting a shared antimicrobial mechanism after prodrug activation. When treating PR strains with Bac, only one PR strain showed visible growth in the presence of 1x MIC Bac, implying the importance of the non-sense mutation of PA14_23460 to the survival and a low likelihood of reduced Bac susceptibility after prodrug exposure (Figure 4C).

## Discussion

Previous protease-activated prodrug designs targeting *Candida albicans* and *Porphyromonas gingivalis* neutralized the cationic nature of the parent AMP by adding a “quencher” sequence with six aspartate residues, which was sufficient for generation of a successfully activated prodrug.^15^ Here we demonstrate that this strategy is unsuccessful in targeting *P. aeruginosa* PA14, and instead a tailored prodrug sequence is required to facilitate activation and avoid potential activity-quenching effects of remaining L-glutamic acid resides in the case of partial activation. Knowledge on PaAP substrate scope was key to the prodrug design, as the pro-moiety had to be tuned for cleavage by specific secreted proteases. We incorporated glutamate and leucine residues in the pro-moiety, as our previous characterization of PaAP determined quick hydrolysis occurred with peptide substrates containing these residues. This tailored prodrug showed accelerated PaAP-mediated activation and in antimicrobial activity in *P. aeruginosa* in planktonic culture and biofilms (Figure 2).

Furthermore, the pro-moiety underwent strain-specific activation in ESKAPEE bacteria, as others lack secreted aminopeptidases with similar substrate scope. Cross activation in mixed cultures with *E. coli* took place, but the population composition affected efficacy of the prodrug, such that excess of competing *E. coli* led to no cross activation.

A similar concept employing bacterial proteases^15,50^ to activate cationic AMP prodrugs has been proposed for *P. aeruginosa*. In that design, the pro-moiety comprised an anionic shielding block (PEG_2k_) and a protease cleavage site (Gly_3_) specific for elastase A, thereby enabling a *P. aeruginosa*-specific antimicrobial activity and improved biosafety towards A549 cells. While the expression of LasA is under the control of LasI/R system, LasI/R-deficient clinical isolates were often isolated from patients, where *P. aeruginosa* are unable to express LasA and LasB, and therefore those may be unable to activate the prodrug *in vivo*.^51^ In the case of ELEG-Bac, PaAP facilitated the activation and contributed to antibiofilm activity of the prodrug in biofilms, but activation did not solely rely on a single bacterial protein in planktonic cultures. This reduces the possibility that *P. aeruginosa* could evade prodrug activation through mutations, as both PaAP and LasB would have to become impaired/absent.

The non-sense mutation of wbpL was previously observed in strains with reduced susceptibility to beta-lactam antibiotics, and this mutation can reduce surface hydrophobicity and facilitate cell aggregation. Our evolutionary study revealed mutations related to O-specific antigen biosynthesis can reduce the antimicrobial susceptibility of Bac and the prodrug to *P. aeruginosa*, which addressed the potential roles of the loss of O-specific antigen to survival in the presence of antimicrobials.

Owing to the overlap in substrate scope of bacterial proteases and the lack of a specific cleavage site in our prodrug, we observed unspecific activation of the prodrug in human serum (Figure 2I). Moreover, antimicrobial activity and biosafety could be examined in a more complex environment, such as animal models with wounds or lung infections. Further investigations would be into the development of the deliver and administration method and testing this strategy with other cationic AMPs showing higher cytotoxicity.

The high manufacturing cost of cationic antimicrobial peptides is a barrier to their application.^5^ Heterologous expression of AMPs has been addressed in producing large quantities of AMPs.^52^ In this system, AMPs are usually fused with partner proteins such as thioredoxin, SUMO, and maltose-binding protein to reduce the toxicity to the host, improve solubility, and facilitate affinity-based purification, after which the active form of AMPs is released by removing the fusion.^53^ Future work could explore heterologous expression of cationic antimicrobial peptides directly in the form of a prodrug, instead of harboring larger tags.

## Conclusion

Antimicrobial peptide prodrugs are promising due to their high selectivity and potentially low tissue toxicity, but most rely on human proteins for activation. We designed a prodrug motif, ELEG, grafted onto the cationic antimicrobial peptide D-Bac8C^Leu2,5^. This prodrug can be activated by *P. aeruginosa* PA14, inhibiting its growth in planktonic culture and biofilms. Reduced susceptibility to Bac and ELEG-Bac was observed, which may be linked to the loss of O-specific antigen on the bacterial surface.

Our work paves the way for utilizing a novel type of activating enzymes for AMP prodrugs. We show that knowledge on substrate preference and kinetics of secreted aminopeptidases/proteases enables the design of a prodrug sequence that extends beyond the addition of a negatively charged sequence to improve activation and activity. Finally, reduced susceptibility to the prodrug does not translate into reduced susceptibility to the parent cationic antimicrobial peptide *in vitro, r*evealing a nuanced picture for the consequences and adaptations to prodrug treatment.

## Supporting information

The Supporting information with protein sequences and raw data for gels, mass spec, cell viability is available free of charge. All raw data shown in the main paper images is available at DOI: 10.6084/m9.figshare.31821499.

## Acknowledgements

Q.G. was funded by Tenovus Research Scotland (Grant: T22-48), C.M.C. was funded by the BBSRC Pathfinder IAA BB/X511183/1 to the University of St Andrews), S.L.S. and the instrument used for protein mass spectrometry was funded by BB/T017686/1. M.D. is funded by Research Ireland (former Science foundation Ireland, SFI) under Grant Numbers 20/COV/8470 and 16/RI/3737 .and RCSI (STaR PhD Programme reference 20270A06). Megan Bergkessel was funded by a UKRI Future Leaders Fellowship ( [MR/T041811/1] and [MR/Z000378/1] ).We thank Jimmy Muldoon (University College Dublin, funded under 18/RI/5702) for the peptide mass spectrometry analyses; Jo Hobbs (University of St Andrews) and the “Enzymology Journal Club” team for insightful discussions on the manuscript.

## Contributions

Manuscript was written by Q.G., but all authors contributed to the final form. All authors have given approval to the final version of the manuscript. Specific contributions are as follows: Q.G. designed and performed experiments, interpreted data, wrote manuscript; M.D. participated in peptide design; A.L., J.R.F.B.C. and N.S.R. synthesized peptides; S.S. and S.L.S. performed high resolution mass spectrometry experiments, analyzed data, revised manuscript; C.M.C. supervised Q.G., C.M.C. and M.D. participated in project conception, analyzed and interpreted data, revised manuscript. M.B. generated microbial strains for co-culture experiments, analyzed and interpreted data, revised manuscript.

## Table of contents graphic

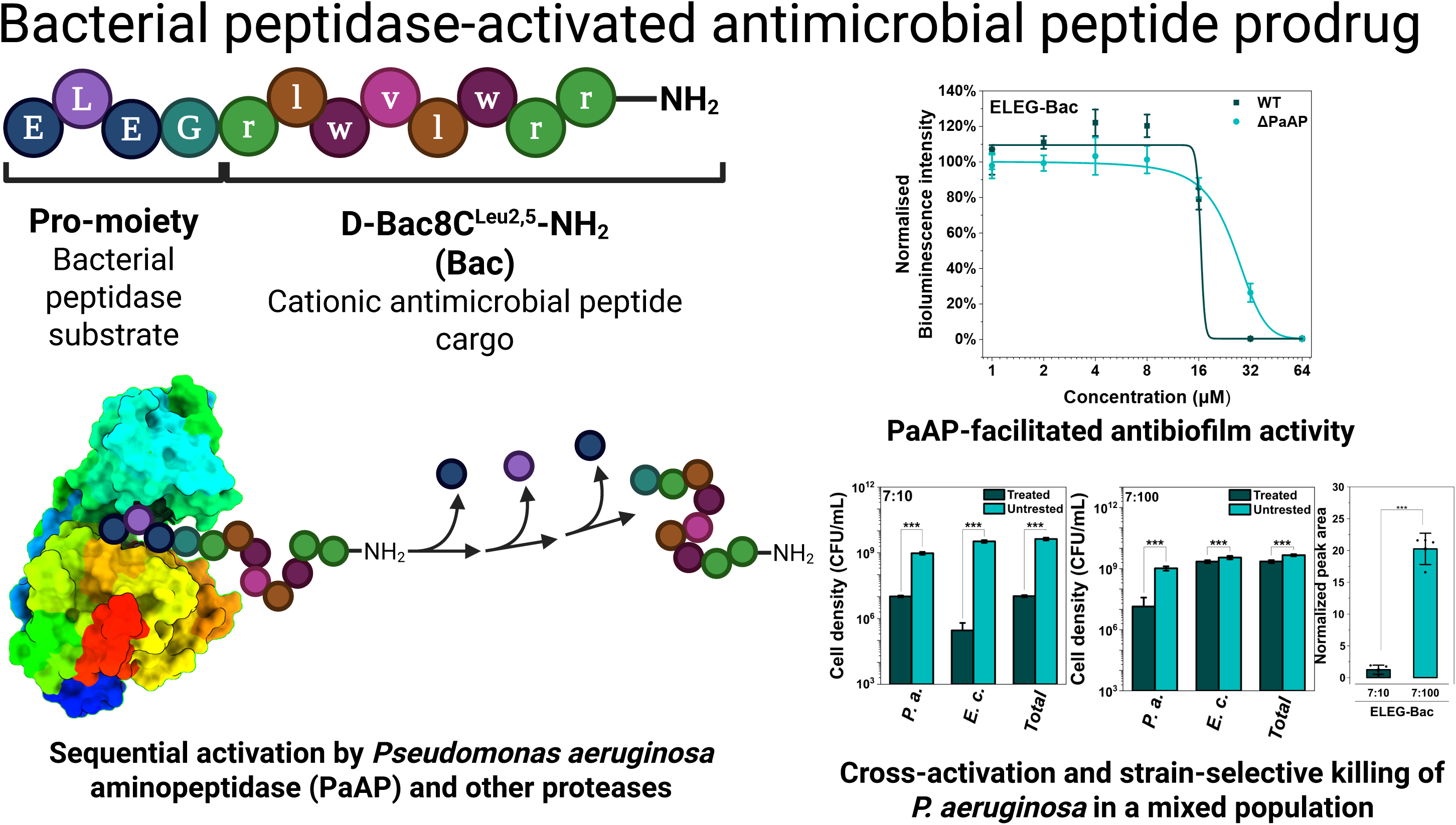

