## Supporting information for "Bacterial Aminopeptidase–Activated Peptide Prodrug Enables Species-Selective Targeting of *Pseudomonas aeruginosa*"

### Supporting Tables

**Table S1.** Primers used

| | |
|---|---|
| CBR_F | GGAATTCGAGCTCAGGAGAAGAAAAAATGGTCAAGCGCGAGAAAAACGTG |
| CBR-mscar_R | GCCTCGCCCTTGCTCACCATGGGATGCACCTATTTTCTAGC |
| CBR-mscar_F | GCTAGAAAATAGGTGCATCCCATGGTGAGCAAGGGCGAGGC |
| mscarlet_R | AGGCCTTCGCGAGGTACCTTACTTGTACAGCTCGTCCAT |
| pUC18_mscar_pF | ATGGACGAGCTGTACAAGTAAGGTACCTCGCGAAGGCCT |
| pUC18_Ptrc_pF | CACGTTTTTCTCGCGCTTGACCATTTTTTCTTCTCCTGAGCTCGAATTCC |
| pMRE_movF | CGACGCGGCCGTGGCGCGCCTCCTAGGTGCTCGAGTCGAGGTCGACGGTATCGATA |
| pMRE_movR | CGGATCCTAGTAAGCCACGTTTTAATTAATCAGGCCTTCGCGAGGTACC |

**Table S2.** DNA sequence of the dsDNA encoding SUMO protein and the pJ411 plasmid DNA used for EMSA assays.

dsDNA:
ATGGGGCACCATCACCATCATCACGCGAGTATGTCAGATTCCGAAGTAAACCAAGAAGCCA
AACCTGAGGTAAAGCCTGAGGTGAAACCGGAAACGCACATTAACTTGAAGGTGTCAGACGG
CTCCTCAGAGATCTTCTTCAAAATCAAAAAGACGACACCGTTGCGTCGCTTAATGGAGGCTTT
CGCAAAGCGTCAAGGTAAAGAGATGGATTCTCTGCGTTTTTTATACGATGGCATCCGTATTCAA
GCTGACCAGACGCCAGAGGATCTTGATATGGAGGATAATGATATTATTGAGGCACATCGCGA
GCAGATTGGCGGT

pJ411 plasmid:

CTACCATCGGCGCTACGGCGTTTCACTTCTGAGTTCGGCATGGGGTCAGGTGGGACCACC
GCGCTACTGCCGCCAGGCAAACAAGGGGTGTTATGAGCCATATTCAGGTATAAATGGGCTC
GCGATAATGTTCAGAATTGGTTAATTGGTTGTAACACTGACCCCTATTTGTTTATTTTTCTAAATAC
ATTCAAATATGTATCCGCTCATGAGACAATAACCCTGATAAATGCTTCAATAATATTGAAAAAGG
AAGAATATGAGCCATATTCAACGGGAAACGTCGAGGCCGCGATTAAATTCCAACATGGATGC
TGATTTATATGGGTATAAATGGGCTCGCGATAATGTCGGGCAATCAGGTGCGACAATCTATCG
CTTGTATGGGAAGCCCGATGCGCCAGAGTTGTTTCTGAAACATGGCAAAGGTAGCGTTGCC
AATGATGTTACAGATGAGATGGTCAGACTAAACTGGCTGACGGAATTTATGCCACTTCCGACC
ATCAAGCATTTTATCCGTACTCCTGATGATGCATGGTTACTCACCACTGCGATCCCCGGAAAA
ACAGCGTTCCAGGTATTAGAAGAATATCCTGATTCAGGTGAAAATATTGTTGATGCGCTGGCA
GTGTTCCTGCGCCGGTTGCACTCGATTCCTGTTTGTAATTGTCCTTTTAACAGCGATCGCGTAT
TTCGCCTCGCTCAGGCGCAATCACGAATGAATAACGGTTTGGTTGATGCGAGTGATTTTGATG
ACGAGCGTAATGGCTGGCCTGTTGAACAAGTCTGGAAAGAAATGCATAAACTTTTGCCATTCT
CACCGGATTCAGTCGTCACTCATGGTGATTTCTCACTTGATAACCTTATTTTTGACGAGGGGAA
ATTAATAGGTTGTATTGATGTTGGACGAGTCGGAATCGCAGACCGATACCAGGATCTTGCCAT

CCTATGGAACTGCCTCGGTGAGTTTTCTCCTTCATTACAGAAACGGCTTTTTCAAAAATATGGT
ATTGATAATCCTGATATGAATAAATTGCAGTTTCATTTGATGCTCGATGAGTTTTTCTAAGCGGCG
CGCCATCGAATGGCGCAAAACCTTTCGCGGTATGGCATGATAGCGCCCGGAAGAGAGTCA
ATTCAGGGTGGTGAATATGAAACCAGTAACGTTATACGATGTCGCAGAGTATGCCGGTGTCTC
TTATCAGACCGTTTCCCGCGTGGTGAACCAGGCCAGCCACGTTTCTGCGAAAACGCGGGA
AAAAGTGGAAGCGGCGATGGCGGAGCTGAATTACATTCCCAACCGCGTGGCACAACAACT
GGCGGGCAAACAGTCGTTGCTGATTGGCGTTGCCACCTCCAGTCTGGCCCTGCACGCGC
CGTCGCAAATTGTCGCGGCGATTAAATCTCGCGCCGATCAACTGGGTGCCAGCGTGGTGGT
GTCGATGGTAGAACGAAGCGGCGTCGAAGCCTGTAAAGCGGCGGTGCACAATCTTCTCGC
GCAACGCGTCAGTGGGCTGATCATTAACTATCCGCTGGATGACCAGGATGCCATTGCTGTG
GAAGCTGCCTGCACTAATGTTCCGGCGTTATTTCTTGATGTCTCTGACCAGACACCCATCAAC
AGTATTATTTTCTCCCATGAGGACGGTACGCGACTGGGCGTGGAGCATCTGGTCGCATTGGG
TCACCAGCAAATCGCGCTGTTAGCGGGCCCATTAAGTTCTGTCTCGGCGCGTCTGCGTCTG
GCTGGCTGGCATAAATATCTCACTCGCAATCAAATTCAGCCGATAGCGGAACGGGAAGGCG
ACTGGAGTGCCATGTCCGGTTTTCAACAAACCATGCAAATGCTGAATGAGGGCATCGTTCCC
ACTGCGATGCTGGTTGCCAACGATCAGATGGCGCTGGGCGCAATGCGCGCCATTACCGAG
TCCGGGCTGCGCGTTGGTGCGGATATCTCGGTAGTGGGATACGACGATACCGAAGATAGCT
CATGTTATATCCCGCCGTTAACCACCATCAAACAGGATTTTCGCCTGCTGGGGCAAACCAG
CGTGGACCGCTTGCTGCAACTCTCTCAGGGCCAGGCGGTGAAGGGCAATCAGCTGTTGCC
AGTCTCACTGGTGAAAAGAAAAACCACCCTGGCGCCCAATACGCAAACCGCCTCTCCCCG
CGCGTTGGCCGATTCATTAATGCAGCTGGCACGACAGGTTTCCCGACTGGAAAGCGGGCA
GTGACTCATGACCAAAATCCCTTAACGTGAGTTACGCGCGCGTCGTTCCACTGAGCGTCAG
ACCCCGTAGAAAAGATCAAAGGATCTTCTTGAGATCCTTTTTTTCTGCGCGTAATCTGCTGCTT
GCAAACAAAAAAACCACCGCTACCAGCGGTGGTTTGTTTGCCGGATCAAGAGCTACCAACT
CTTTTTCCGAAGGTAACTGGCTTCAGCAGAGCGCAGATACCAAATACTGTTCTTCTAGTGTAG
CCGTAGTTAGCCCACCACTTCAAGAACTCTGTAGCACCGCCTACATACCTCGCTCTGCTAAT
CCTGTTACCAGTGGCTGCTGCCAGTGGCGATAAGTCGTGTCTTACCGGGTTGGACTCAAGA
CGATAGTTACCGGATAAGGCGCAGCGGTCGGGCTGAACGGGGGGTTCGTGCACACAGCC
CAGCTTGGAGCGAACGACCTACACCGAACTGAGATACCTACAGCGTGAGCTATGAGAAAGC
GCCACGCTTCCCGAAGGGAGAAAGGCGGACAGGTATCCGGTAAGCGGCAGGGTCGGAA
CAGGAGAGCGCACGAGGGAGCTTCCAGGGGGAAACGCCTGGTATCTTTATAGTCCTGTCG
GGTTTCGCCACCTCTGACTTGAGCGTCGATTTTTGTGATGCTCGTCAGGGGGGCGGAGCCT
ATGGAAAAACGCCAGCAACGCGGCCTTTTTACGGTTCCTGGCCTTTTGCTGGCCTTTTGCTC
ACATGTTCTTTCCTGCGTTATCCCCTGATTCTGTGGATAACCGTATTACCGCCTTTGAGTGAGC
TGATACCGCTCGCCGCAGCCGAACGACCGAGCGCAGCGAGTCAGTGAGCGAGGAAGCG
GAAGGCGAGAGTAGGGAACTGCCAGGCATCAAACTAAGCAGAAGGCCCCTGACGGATGG
CCTTTTTGCGTTTCTACAAACTCTTTCTGTGTTGTAAAACGACGGCCAGTCTTAAGCTCGGGC
CCCCTGGGCGGTTCTGATAACGAGTAATCGTTAATCCGCAAATAACGTAAAAACCCGCTTCG
GCGGGTTTTTTTATGGGGGGAGTTTAGGGAAAGAGCATTTGTCAGAATATTTAAGGGCGCCTG
TCACTTTGCTTGATATATGAGAATTATTTAACCTTATAAATGAGAAAAAAGCAACGCACTTTAAAT
AAGATACGTTGCTTTTTCGATTGATGAACACCTATAATTAAACTATTCATCTATTATTTATGATTTTTT

GTATATACAATATTTCTAGTTTGTTAAAGAGAATTAAGAAAATAAATCTCGAAAATAATAAAGGGAA
AATCAGTTTTTGATATCAAAATTATACATGTCAACGATAATACAAAATATAATACAAACTATAAGAT
GTTATCAGTATTTATTATGCATTTAGAATAAATTTTGTGTCGCCCTTCCGCGAAATTAATACGACT
CACTATAGGGGAATTGTGAGCGGATAACAATTCCCCTCTAGAAATAATTTTGTTTAACTTTTAAG
GAGGTAAACATATGCACCATCACCATCATCACGACTATGATATTCCCACAACTGAAAATTTGTA
TTTCCAGGGGATGGCGACGGTTTTAATTCAAGGAGCTGGTATTGCTGGACTGGCCTTAGCCC
GCGAGTTCACCAAGGCGGGGATTGATTGGCTGCTTGTTGAGCGTGCGTCAGAGATTCGTCC
TATCGGGACTGGTATCACCCTGGCTTCCAACGCCCTTACCGCATTGTCCAGTACGCTGGATC
TGGATCGCTTATTCCGTCGCGGAATGCCGCTGGCCGGAATTAATGTGTATGCACACGACGG
GTCGATGTTAATGTCCATGCCGAGCAGCCTTGGTGGGTCAAGCCGCGGTGGCTTGGCCTTG
CAGCGCCACGAGCTTCATGCTGCGTTGTTGGAGGGTTTAGACGAGAGCCGCATTCGTGTGG
GTGTCAGTATTGTACAGATTCTTGATGGGCTGGATCACGAACGCGTGACGTTATCCGATGGTA
CCGTTCACGACTGTAGTCTTGTGGTGGGCGCTGACGGCATTCGTTCTAGTGTGCGTCGCTAC
GTCTGGCCTGAGGCTACGTTACGTCACTCTGGTGAAACTTGCTGGCGTTTAGTAGTCCCTCA
CCGCTTGGAAGATGCCGAGTTAGCCGGAGAAGTTTGGGGTCATGGAAAACGTCTGGGCTTC
ATTCAGATTAGTCCTCGTGAAATGTACGTTTATGCTACCTTAAAAGTGCGCCGTGAAGAGCCC
GAGGATGAGGAAGGTTTTGTAACTCCACAGCGTTTGGCTGCCCATTATGCAGATTTCGACGG
AATCGGTGCTTCAATTGCGCGCTTAATCCCTTCTGCAACTACTCTGGTCCACAATGATCTGGA
AGAACTTGCAGGTGCCTCCTGGTGTCGTGGACGTGTGGTGCTGATCGGGGACGCGGCGCA
CGCGATGACCCCGAACCTGGGACAAGGAGCGGCGATGGCCTTGGAGGACGCGTTCCTTT
TAGCTCGCTTATGGTGCTTAGCCCCACGTGCTGAGACATTAATTCTGTTTCAACAACAGCGCG
AAGCCCGCATTGAATTTATCCGTAAGCAGAGTTGGATTGTCGGACGTTTAGGTCAATGGGAGA
GCCCTTGGTCGGTTTGGCTGCGCAACACATTGGTCCGTCTGGTCCCGAATGCCAGCCGTC
GTCGCTTACATCAGCGCTTGTTCACGGGGGTTGGAGAGATGGCGGCGCAATGAAAGCTTCC
CCCTAGCATAACCCCTTGGGGCCTCTAAACGGGTCTTGAGGGGTTTTTTGCCCCTGAGACG
CGTCAATCGAGTTCGTACCTAAGGGCGACACCCCCTAATTAGCCCGGGCGAAAGGCCCAG
TCTTTCGACTGAGCCTTTCGTTTTATTTGATGCCTGGCAGTTCCCTACTCTCGCATGGGGAGT
CCCCACA

**Table S3.** Retention times and the m/z for selected ions in LC-MS experiments.

| Ion | m/z (Number of charges) | LC column | Retention time (min) |
|---|---|---|---|
| EEEE-D-Bac8C | 850.97 (+2) | Kinetex XB-C18 2.6 µm 100 Å | 6.82 |
| EEE-D-Bac8C | 786.41 (+2) | | 6.80 |
| EE-D-Bac8C | 721.85 (+2) | | 6.80 |
| E-D-Bac8C | 657.29 (+2) | | 6.78 |
| D-Bac8C | 592.73 (+2) | | 6.70 |
| EEEE-D-Bac8C | 850.97 (+2) | XSelect PREMIER HSS T3 2.5 µm 4.6 x 50 mm 100 Å | 7.00 |
| EEE-D-Bac8C | 786.41 (+2) | | 7.00 |
| EE-D-Bac8C | 721.85 (+2) | | N.D. |
| E-D-Bac8C | 657.29 (+2) | | N.D. |
| D-Bac8C | 592.73 (+2) | | 7.85 |
| ELEG-D-Bac8C | 538.30 (+3), 403.73 (+4) | | 8.28 |
| LEG-D-Bac8C | 495.29 (+3), 371.47 (+4) | | 8.15 |
| EG-D-Bac8C | 457.60 (+3), 343.20 (+4) | | 8.00 |
| G-D-Bac8C | 414.60 (+3) 310.94 (+4) | | 7.78 |

**Table S4.** Parameters for fitted curves in Figure 2A.

| Curve fitting of the first 240 mins | | | | | | | | | | |
|---|---|---|---|---|---|---|---|---|---|---|
| | Peptide | $y_0$ | Error | $k_1$ | Error | Model | $k_2$ | Error | Rate of linear phase | Error |
| ELEG | ELEGrlwvlwrr-$NH_2$ | 13.39 | 0.63 | 0.030 | 0.001 | Single exponential | | | | |
| LEG[a] | LEGrlwvlwrr-$NH_2$ | 1.47 | 0.04 | Very fast | | Single exponential for phase 2 | 0.032 | 0.001 | | |
| EG | EGrlwvlwrr-$NH_2$ | 12.69 | 2.86 | 0.021 | 0.006 | Single exponential+ Liner phase | | | 0.042 | 0.017 |
| G | Grlwvlwrr-$NH_2$ | | | | | Lag phase+ Linear phase | | | 0.035 | 0.001 |

[a] Rate constant $k_1$ and $y_0$ of the first exponential phase cannot be fitted reliably due to insufficient information of the amplitude. $y_0$ shown in the table is from the curve fitting of data from 15 minute to the end of the experiment.

**Table S5.** Parameters for fitted curves in Figure 2H.

| Strain | Treatment | $y_0$ | Error | $t_1$ | Error | Model | Rate of linear phase | Error |
|---|---|---|---|---|---|---|---|---|
| WT | Blank | | | | | Linear | 0.843 | 0.220 |
| WT | Bac | 2137.941 | 14.218 | 49.085 | 2.797 | Single exponential | | |
| WT | ELEG-Bac | 2804.820 | 247.068 | 106.967 | 23.819 | Single exponential+ Linear phase | 5.812 (t0=141.807±10.264) | 1.004 |
| ΔPaAP | Blank | | | | | Linear | 1.013 | 0.163 |
| ΔPaAP | Bac | -1688.476 | 35.404 | 45.952 | 2.271 | Single exponential | | |
| ΔPaAP | ELEG-Bac | | | | | Linear | 3.525 | 0.149 |

### Supporting Figures

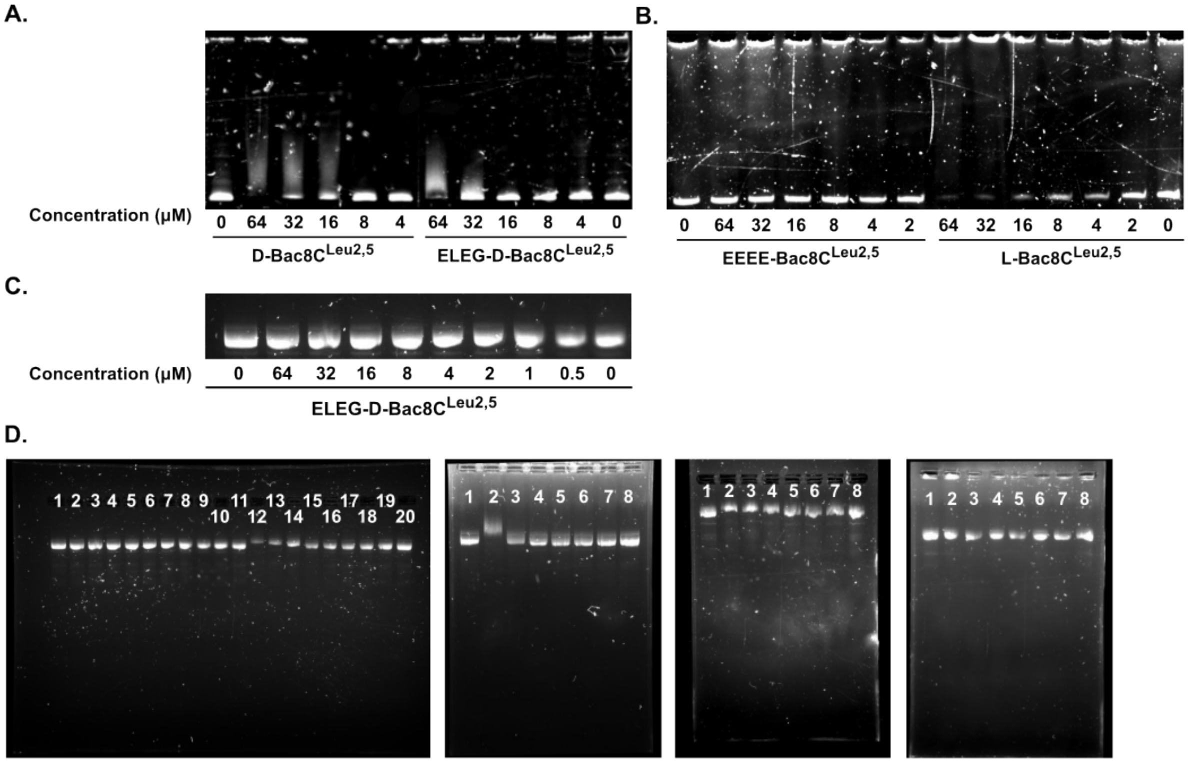

**Figure S1.** Electrophoretic mobility shift assay (EMSA) showing the formation of DNA-AMP complex. (A) Binding of a 326-bp double strained DNA with D-Bac8CLeu2,5 and ELEG-Bac; (B) binding of the 326-bp double strained DNA with EEEE-Bac and L-Bac8C$^{Leu2,5}$; (C) binding of the pJ411 plasmid DNA with ELEG-Bac; (D) raw gel images from Figure 1. From left to right, 1st gel: interactions of the pJ411 plasmid DNA to ELEG-Bac (Lane 1-11) and Bac (Lane 12-20); 2nd gel: interaction between the pJ411 plasmid DNA to L-Bac8CLeu2,5; 3rd gel: interaction of the pJ411 plasmid DNA to Bac in the presence of a concentration series of NaCl; 4th gel: interaction of the pJ411 plasmid DNA to EEEE-Bac.

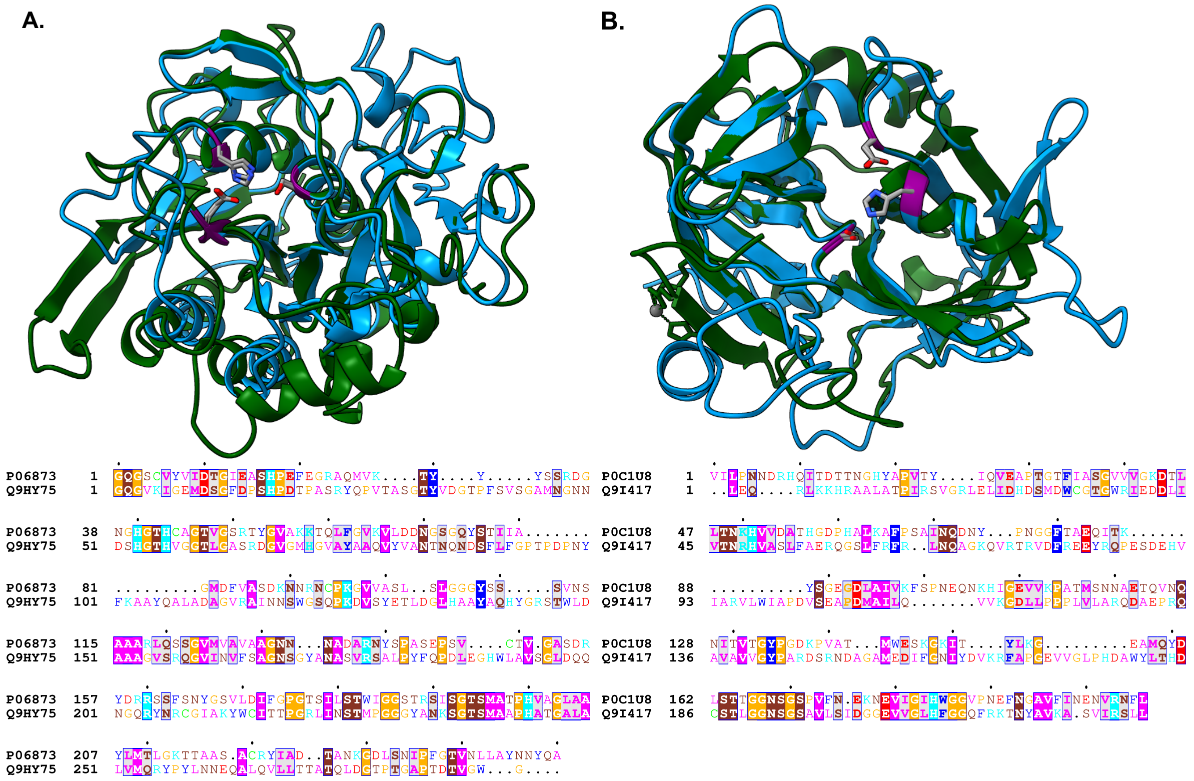

**Figure S2.** Structure alignment indicated the potential activities of two proteins in P. aeruginosa PA14 genome. Sequence alignment of Proteinase K from Tritirachium album (UniProtKB ID: P06873) (A) and staphylococcal peptidase I (SspA) from S. aureus (UniProtKB ID: P0C1U8) (B) with the peptidase domain from the homologue in P. aeruginosa. Catalytic triads are highlighted in purple with atoms. Proteinase K (PDB ID: 2PKC) and SspA (PDB ID: 1WCZ) were reference structures and were colored in dark green. The peptidase domain of PA14_47090 (residue 107-330, Q9HY75) and PA14_18630 (residue 70-359, Q9I417) were identified by InterPro and renumbered in the sequence alignment.

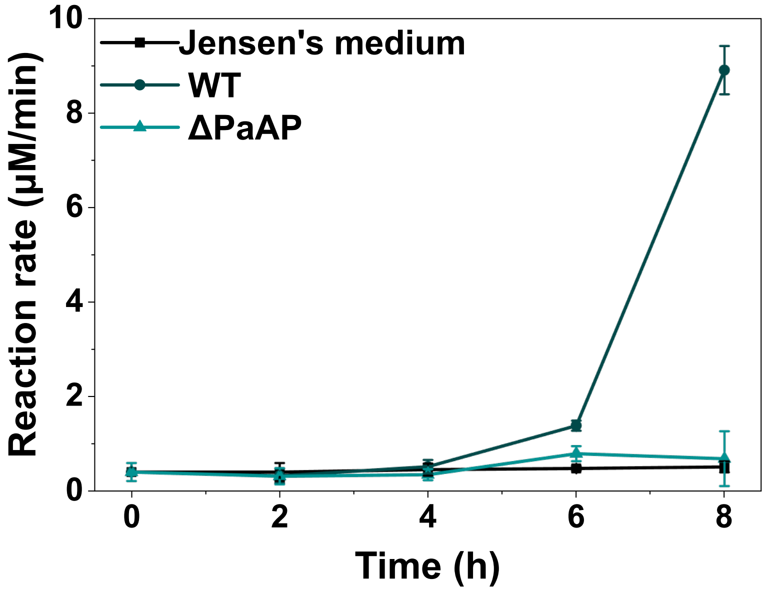

**Figure S3.** Leucine aminopeptidase activity from *P. aeruginosa* WT culture in Jensen's medium.

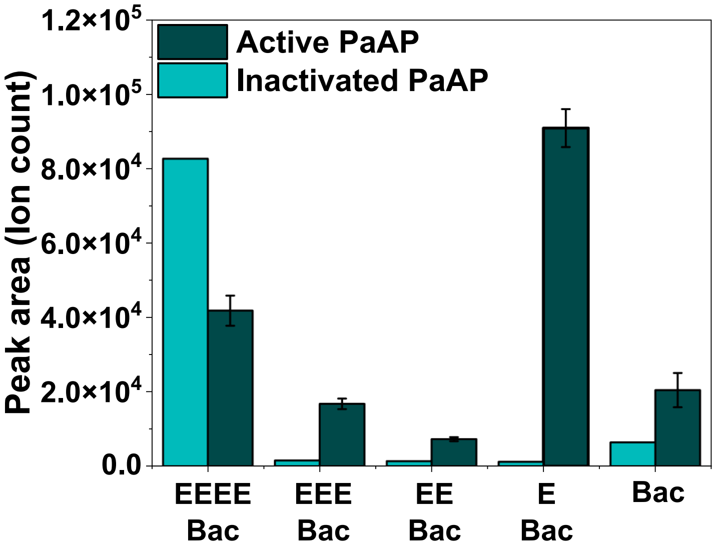

**Figure S4.** Relative quantification of each intermediate of EEEE-Bac in an in vitro PaAP activation experiment.

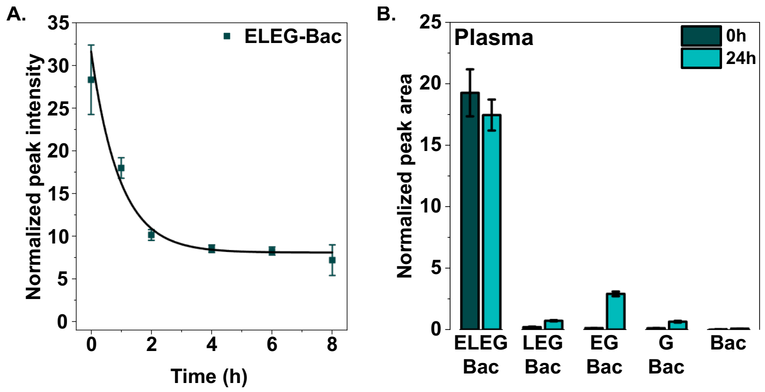

**Figure S5.** Prodrug stability in human serum and human plasma. (A) Relative quantification of ELEG-Bac in human serum during 8-hour incubation at 37℃; (B) relative quantification of ELEG-Bac in human plasma after 24-hour incubation at 37℃.

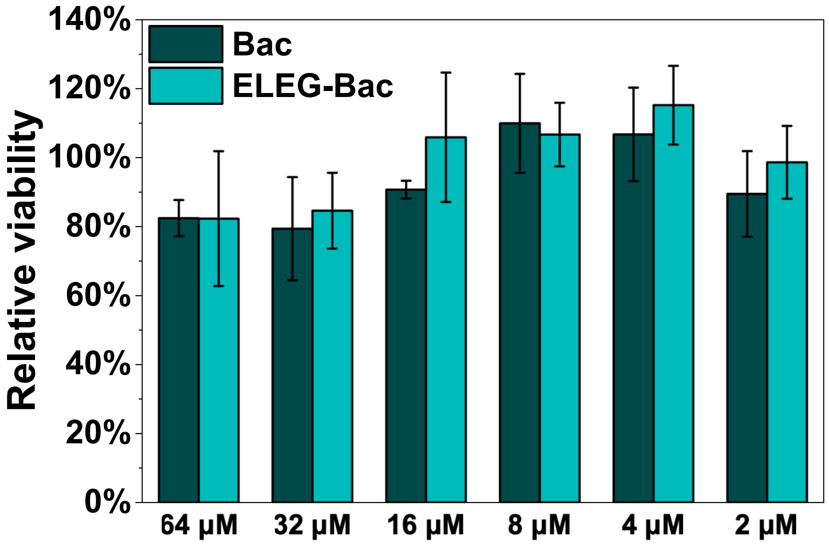

**Figure S6.** Viability of HEK293FT cells after 24-hour treatment of Bac or ELEG-Bac.

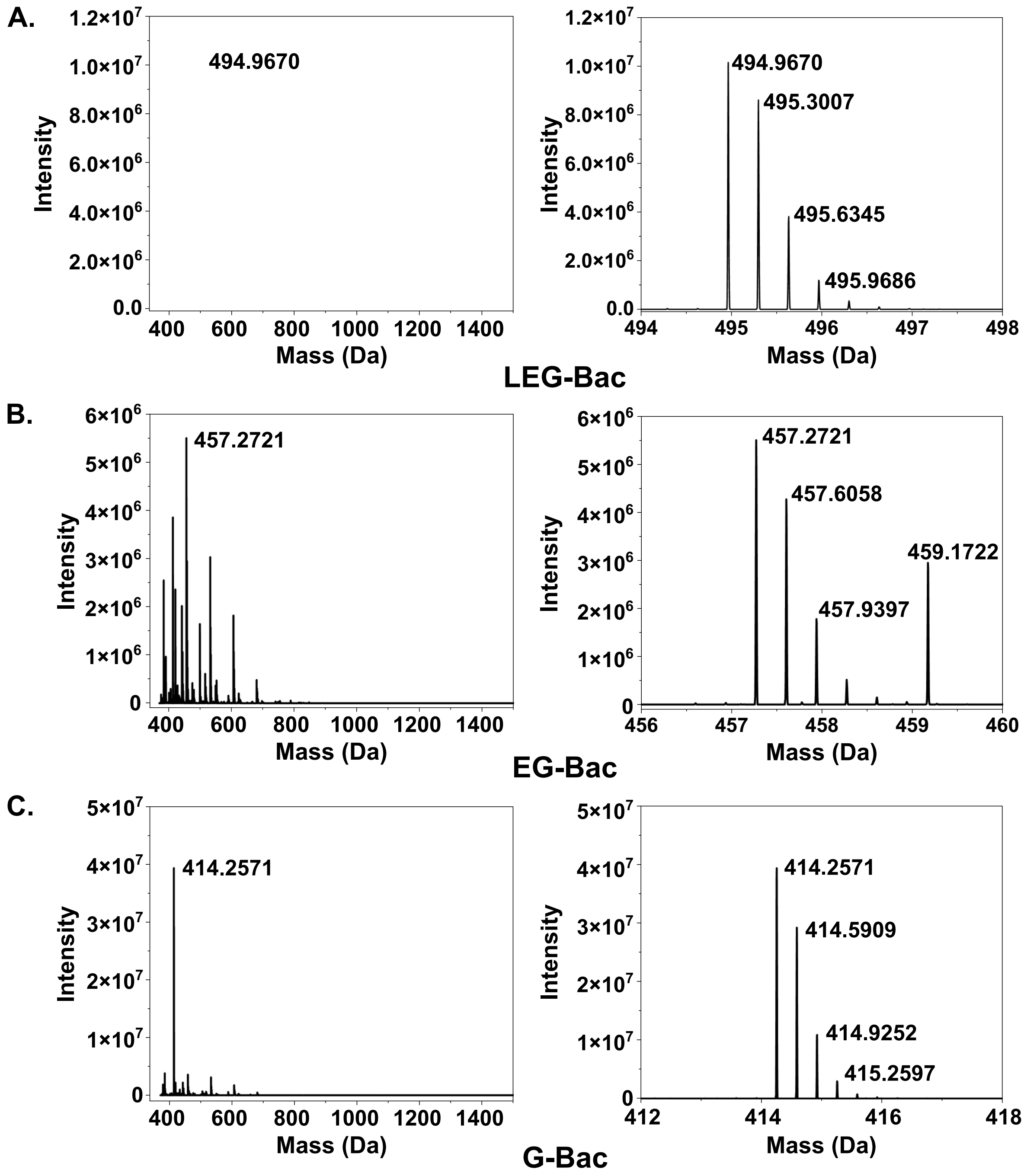

**Figure S7.** MS spectra of the elution peak for LEG-Bac (A), EG-Bac (B), and G-Bac (C) from the bacteria culture with 32 μM ELEG-Bac after the experiment in Figure 2B and 2C. The full spectra of each elution peak (top) and the zoom-in spectra of the peak with the highest intensity (bottom) were illustrated.

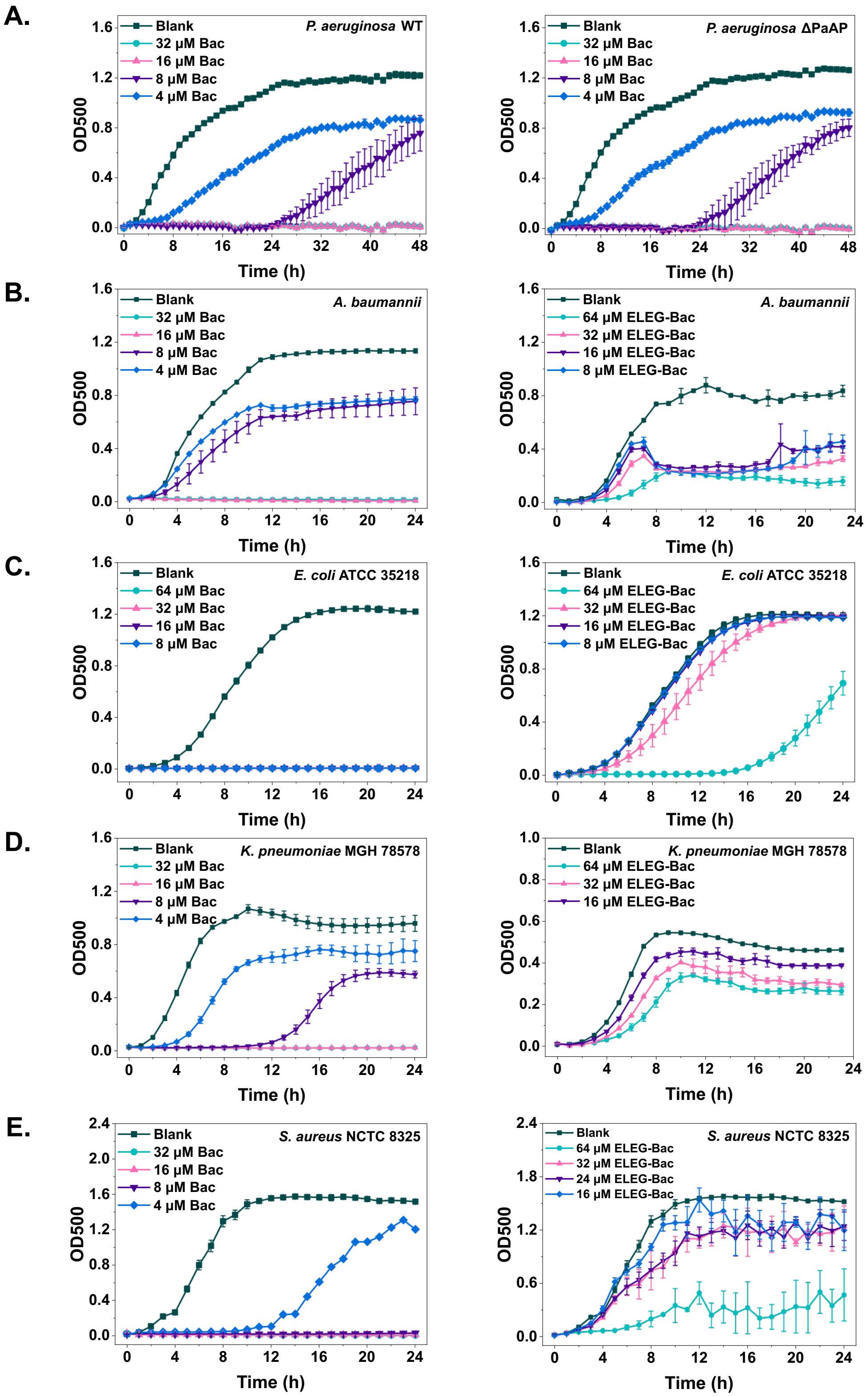

**Figure S8.** Antimicrobial tests of bacteria in the presence of Bac and ELEG-Bac. Growth curves of (A) two P. aeruginosa strains with Bac; (B) A. baumannii; (C) E. coli ATCC 35218; (D) K. pneumoniae MGH 78578; (E) S. aureus NCTC 8325 with Bac and ELEG-Bac.

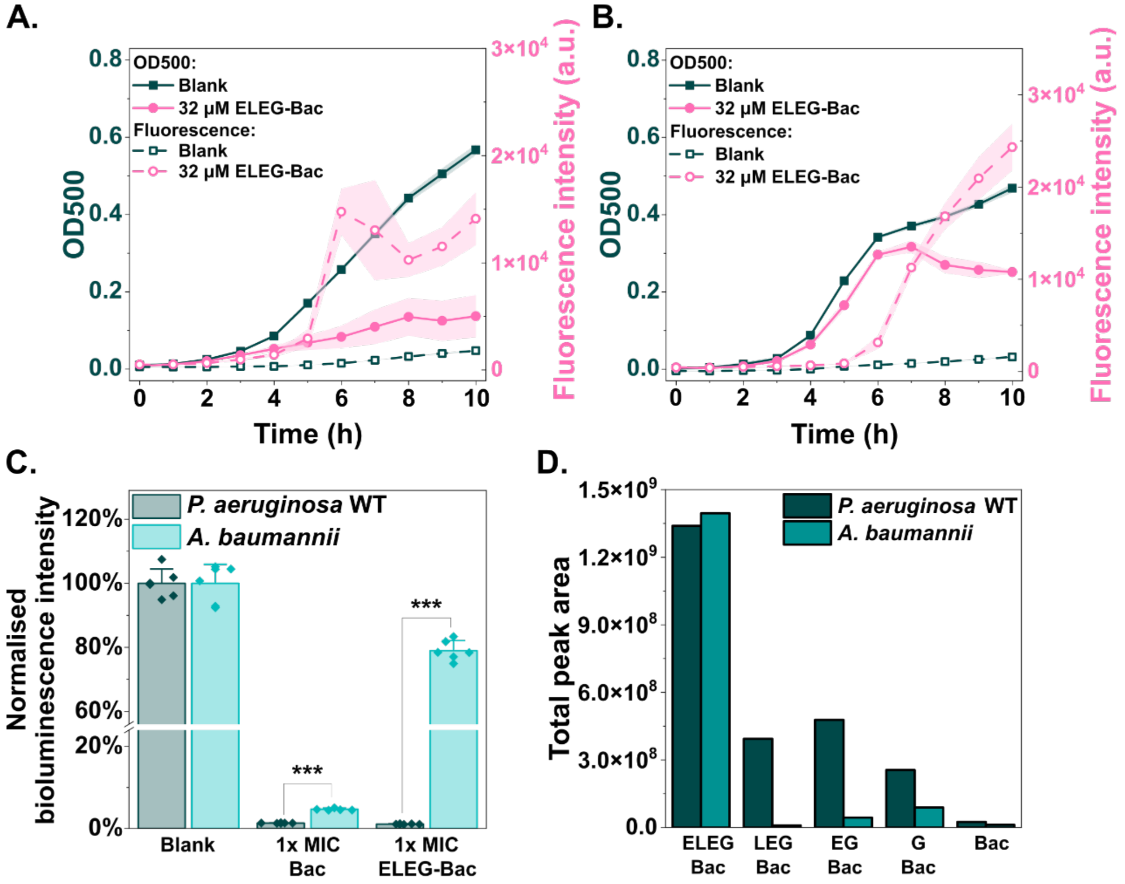

**Figure S9.** The prodrug influenced the growth of *A. baumannii* despite limited activation. (A-B) Growth curves (dark green solid symbols with solid lines at the left Y-axis) and PI fluorescence (pink hollow symbols with dash lines at the right Y-axis) of (A) P. aeruginosa and (B) A. baumannii cultures with 32 μM ELEG-Bac; (C) total metabolic activity after experiments in Figure 3B&C; (D) quantification of intermediates in the whole cell culture after experiments for panel A and B. (*: $p<0.05$; **: $p<0.01$; ***: $p<0.001$)

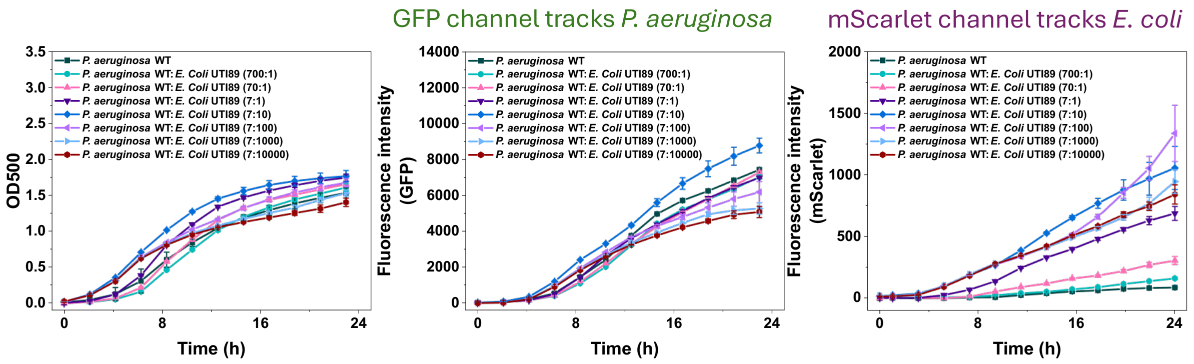

**Figure S10.** Growth of GFP-labelled P. aeruginosa WT strain and mScarlet-labelled E. coli strain at the mixing ratio from 700:1 to 7:10000 (cell count). Panels from left to right are growth curves in OD, fluorescence intensity tracking GFP-labelled P. aeruginosa WT strain, and fluorescence intensity tracking mScarlet-labelled E. coli UTI89 strain.

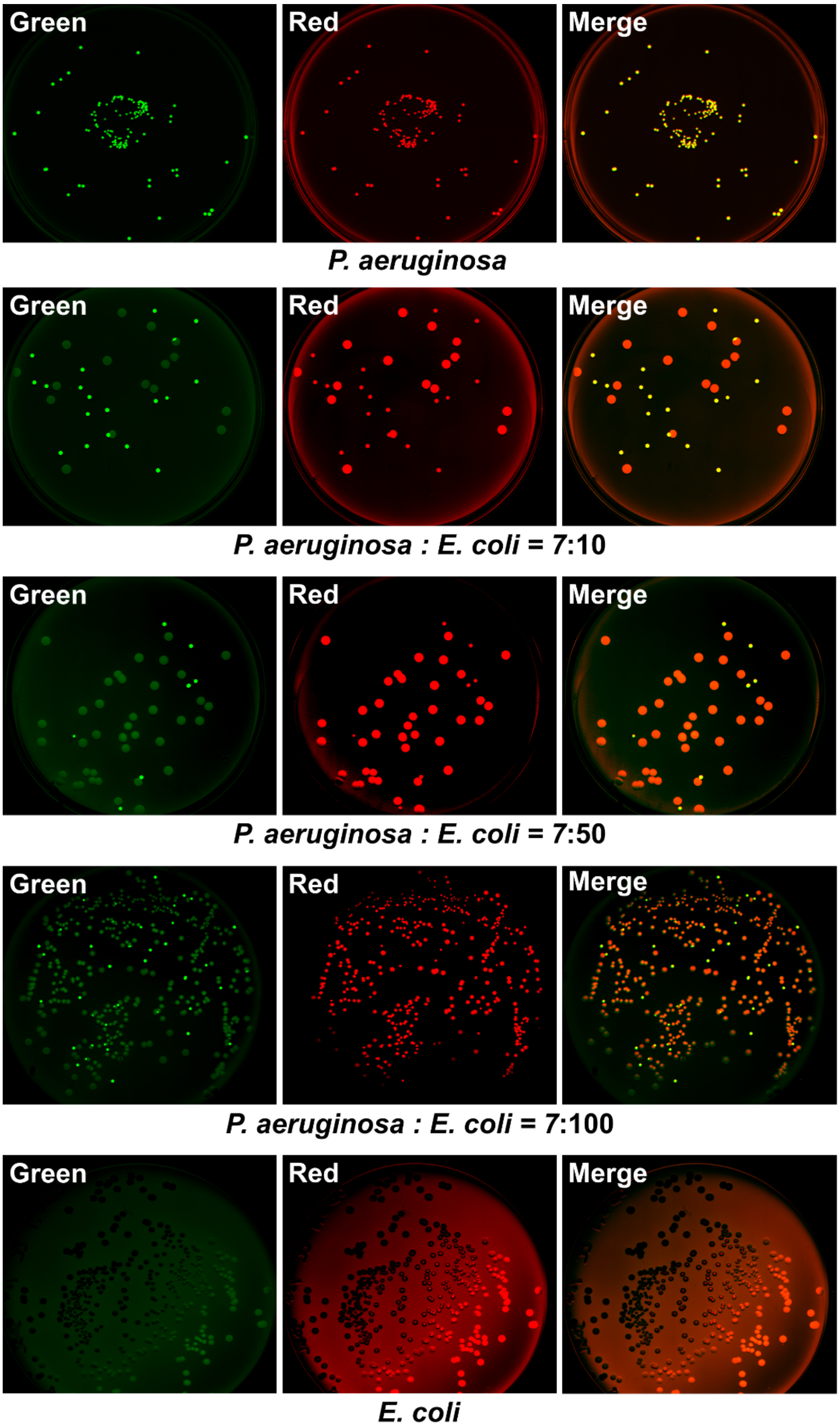

**Figure S11.** LB agar images for identification and quantification of GFP-tagged *P. aeruginosa* WT strain and mScarlet-tagged *E. coli* UTI89 strain by fluorescence at a certain mixing ratio. Cell mixture was diluted 5x104 times, and 50 µL was used to inoculate LB agar plates. Fluorescence intensity was adjusted in LI-COR acquisition software to highlight the edge of LB agar plates.

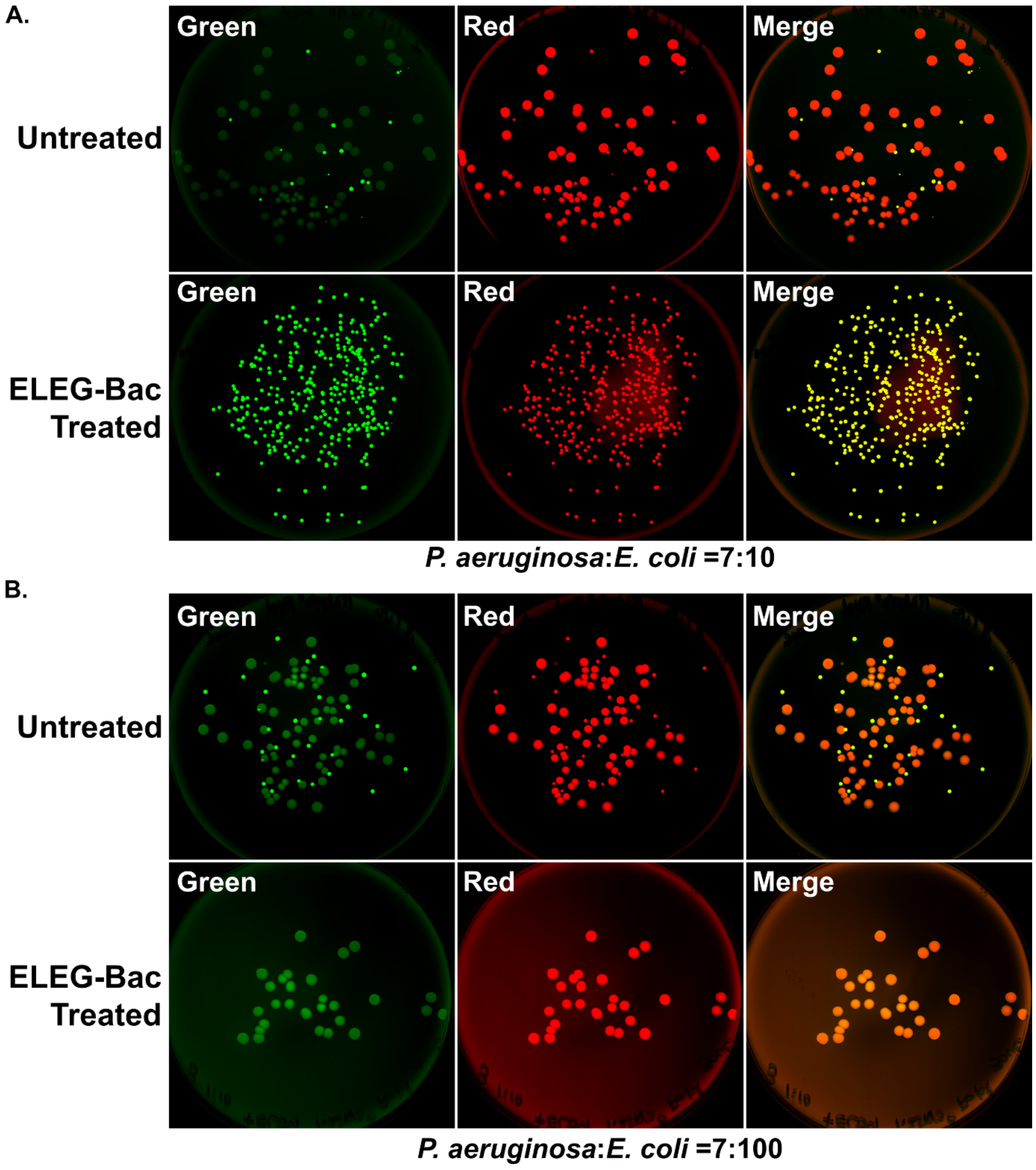

**Figure S12.** Representative LB agar images for quantification of GFP-tagged P. aeruginosa WT strain and mScarlet-tagged E. coli UTI89 strain by fluorescence at the end of experiments for Figure 3B (Figure S12A) and 3C (Figure S12B). Mixed culture in the absence (Untreated) and presence of 50 μM ELEG-Bac (ELEG-Bac treated) was diluted 1.25x106- and 1000-fold, respectively. 50 μL was used to inoculate LB agar plates. Images in green, red, and their merged channel were listed from left to right. All experiments were carried out with three biological replicates.

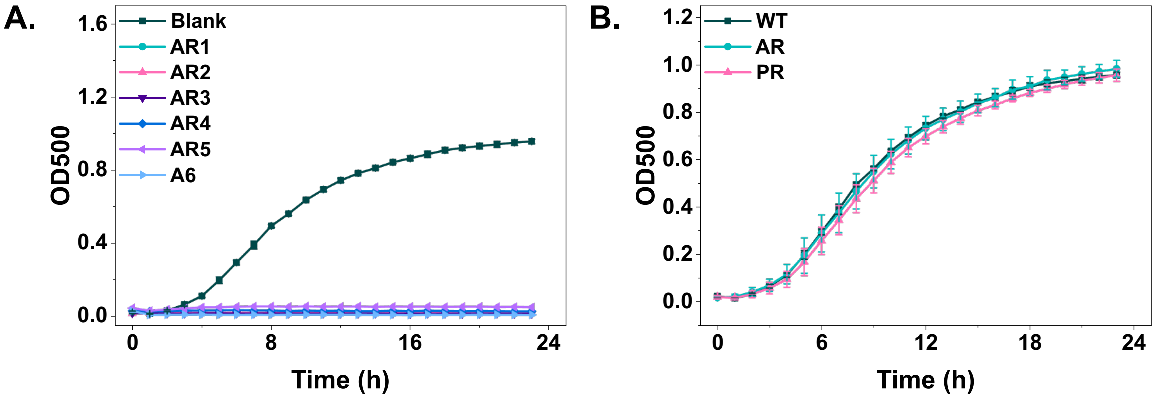

**Figure S13.** Evaluation of (A) antimicrobial susceptibility of AR strains with 32 μM Bac. Blank is a growth curve in the absence of inhibitor. (B) depicts lack of fitness cost for all AR and PR strains.

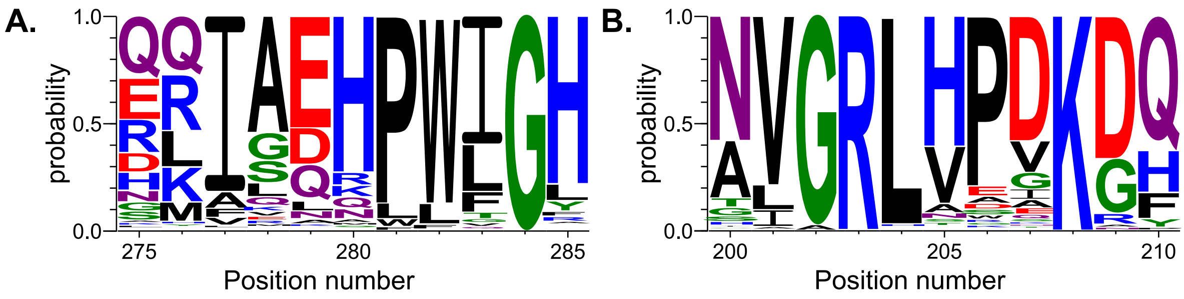

**Figure S14.** Sequence alignment revealed that L204P and H280P mutations are not frequently present in homologues from other species. WebLogo represents the sequence alignment of (A) PA14_66100 and (B) PA14_66160 relative to homologues.
